# A validation approach for RNA structures based on deep-learning 3D-structure predictions

**DOI:** 10.64898/2026.11.02.763582

**Authors:** Aderik Voorspoels, Adam J Simpkin, Ronan M Keegan, Grzegorz Chojnowski, Daniel J Rigden

## Abstract

Understanding the molecular mechanisms of life depends on accurate macromolecular structures. RNA, in particular, poses a particular challenge for structural biology due to its inherent flexibility and dynamic nature. The challenging nature of structure determination experiments and their interpretation have led to the development of validation approaches to prevent or fix errors in solved structures.

Existing methods for RNA structure validation can, as with proteins, be broadly split into two types. Map-based validation tools seek to detect discrepancies between the experimental data and the determined structure. They are excellent for fixing all types of errors, but only when high-resolution experimental data are available. Model-only validation methods, which usually focus on molecular geometry, are excellent for fixing short-lengthscale issues even when experimental resolution is low. *Conkit-validate* is a mapless method for protein validation, which is not based on chemical plausibility, but rather on the similarity between experimental models and structure predictions for the same protein. By comparing contact patterns, this tool leverages evolutionary covariance information, learned by state-of-the-art structural prediction models, into a mapless validation method capable of detecting sequence assignment issues, with notable success in detecting register errors.

Here, we extend the *conkit-validate* method to work on RNA. The results show performance similar to that previously achieved for proteins. In a scan of the PDB, its detection rate does not depend on the local resolution of the density, so it remains applicable where map-based approaches are not, and case studies show it can identify register errors where the density alone cannot decide between models. Together, this highlights the complementary value of a contact-prediction based validation approach using information orthogonal to map- and geometry-based approaches to ensure the quality of deposited RNA structures against varying criteria. The performance of the method can be expected to track ongoing improvements in RNA structure prediction methods.

## Introduction

Structured RNA plays a crucial role in biology. Correspondingly, the determination and study of high quality RNA structures plays a crucial role in structural biology. Detailed and accurate knowledge of RNA 3D structure is obtained using experimental techniques such as X-ray crystallography and cryo-EM. Each method provides a map, of the electron density or Coulomb potential respectively, leaving the interpretation of those maps as detailed atomic structures to the experimentalist. While great arsenals of tools (1) are available for this task of interpretation, errors are known to occur in the process. These can be the result of low resolution experimental information or reliance on faulty pre-existing structures. Unfortunately such errors do occasionally end up deposited in the Protein Data Bank (PDB)(2), with lower resolution cryo-EM structures posing special concern(3).

Validation tools continue to be developed to detect and mitigate the impact of these errors. These include methods that operate based on the underlying experimental data or map (4, 5). Such tools are great tests of how faithful the interpretation is to the actual experiment, but they suffer from applicability issues at lower resolutions where errors are most common. A different class of validation tools works on a model-only basis but instead focuses on the internal consistency of the structural model. These methods tend to validate the local geometry of the structure by checking if the backbone (6) conforms to known geometrical classes, by checking for deviations in bond angles and lengths(7), or by checking for clashes(8). While those methods can directly improve the quality of structure on a local level, they are blind to more global problems such as register shift and sequence assignment errors.

In the protein world, one tool that addresses this precise problem of model-only validation for register-related errors is *conkit-validate*(9). It works by comparing a candidate structure against coevolutionary information, as summarised by structure prediction methods, employing both machine learning and classical computation approaches to do so. Rather than straightforwardly using predicted structures, the method is built around predicted distograms which form part of the structure prediction pipeline. These were preferred as input over predicted coordinates, as they are able to encode coevolutionary contact information more directly without also encoding a specific conformation. A scan of the PDB using this tool revealed that, at least for proteins, a non-trivial number of structures suffer from register shift errors(10).

The development of the original *conkit-validate* tool was enabled by the leap in accuracy of protein structure prediction methods initiated with *AlphaFold 2*(11). Since then, the capabilities of similar structure prediction methods have expanded beyond single protein chains to include protein complexes and most relevantly nucleic acids. This has led to a broad generation of distinct but similar prediction models (12–14) finding applications in many areas of structural biology. While critical assessment (CASP)(15) and specialised RNA folding reviews(16) found automated structure predictions of RNA lagged behind proteins(17) and are still outperformed by human experts, RNA-specific methods continue to improve (18, 19).

As RNA structure prediction continues to become more reliable and accessible, the extension of *conkit-validate* to RNA has become feasible. In this publication we introduce a validation method for RNA structures based on coevolutionary information as summarised by structure or distogram predictions (Fig. 1). To do this, first a set of known errors to benchmark the methods was created. Then necessary concepts like a base level contact definition were transposed to RNA. Subsequently the contact map overlap method was benchmarked for RNA and a machine learning classifier was trained to detect errors in RNA. Finally a subset of PDB structures was scanned with the new method and some useful case studies are described in detail.

**Fig. 1.**
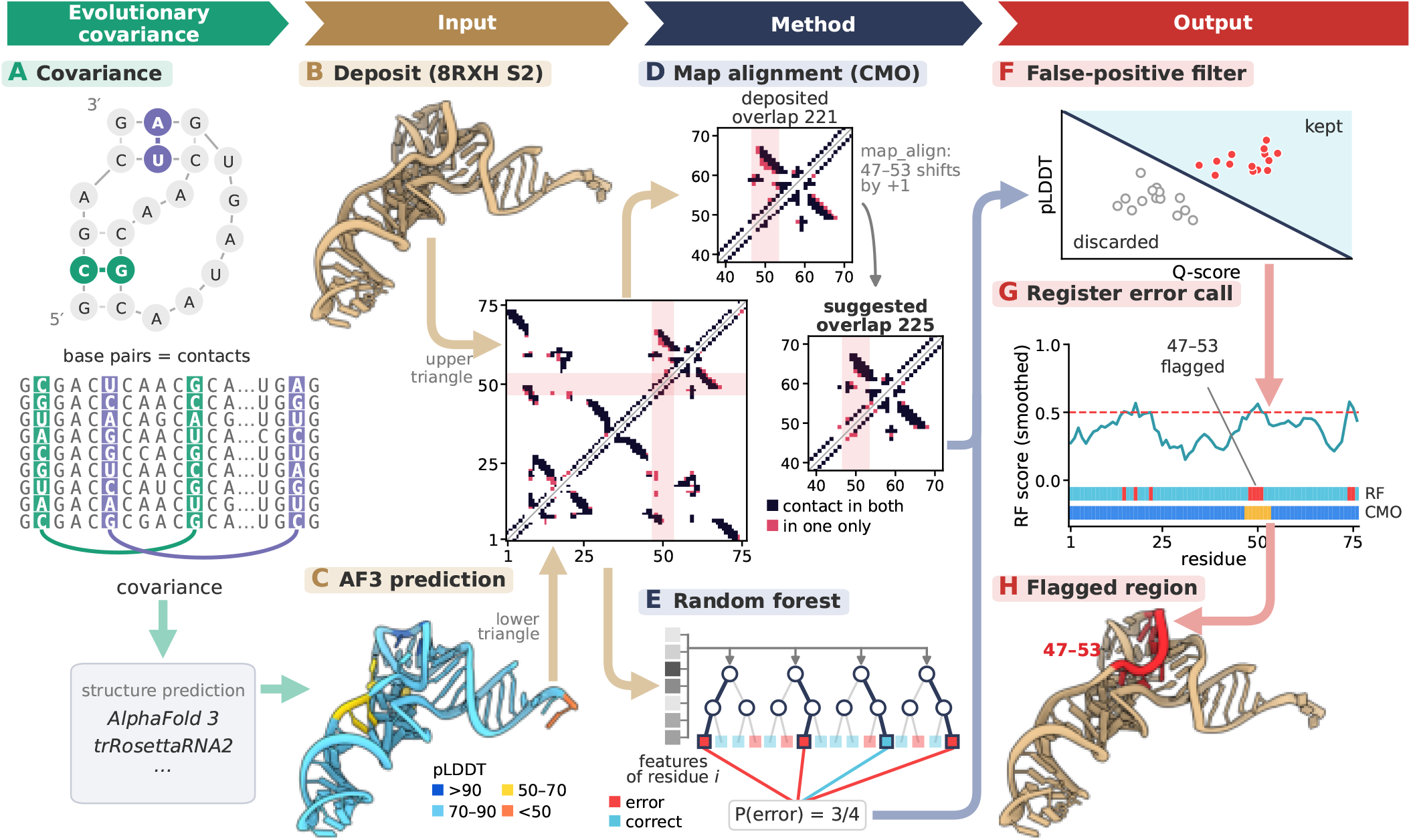
Operation of the *conkit-validate* pipeline, shown on 8RXH chain S2. (A) Covarying base pairs in an alignment of homologous sequences (schematic) are the evolutionary signal that structure predictors such as *AlphaFold 3* and *trRosettaRNA2* draw on. (B, C) The deposited chain and its AF3 prediction (coloured by pLDDT) are each turned into a contact map (C1^*′*^–C1^*′*^ distance *≤* 12.5 Å, covalent neighbours excluded), with the deposit above and the prediction below the diagonal; contacts present in both are shown dark, contacts present in only one structure in pink. (D) Contact map alignment (CMO) with *map_align* re-registers the deposit to maximise its overlap with the prediction; for 8RXH S2, shifting residues 47–53 by +1 raises the overlap from 221 to 225 contacts. (E) A random forest classifier (schematic) assigns each residue a probability of being in error from per-residue features comparing the two contact maps. (F) Calls from both routes pass a LASSO false-positive filter (schematic) on metrics not used to make them, such as pLDDT and Q-score. (G) The output for 8RXH S2: the smoothed random forest score with its threshold, and the residues flagged by the random forest (RF) and CMO, here 47–53. (H) The flagged region on the deposited structure. Panels A, E and F are schematics; all other panels show the data for 8RXH S2.

## Methods

### Benchmark dataset creation

To create a set of known modelling errors, all single chain RNA structures of less than 1000 residues, solved by cryo-EM at a resolution better than 5 Å, released in the PDB before the 8th of May 2025, with a corresponding reconstruction available in EMDB were selected. This resulted in a set of 57 structures. As a scan with *checkMySequence* (CMS)(20), or more specifically its nucleic acid extension *DoubleHelix*(4), revealed no errors in these structures, they were considered to represent a groundtruth set.

Subsequently, the Coulomb potential maps corresponding to the ground truth set were obtained from EMDB. To these maps a low pass filter was applied using *ChimeraX*(21) (setting sDev 1 and step 1), to obtain a set of lower resolution versions of the maps. *DeepCryoRNA*(22) was then used to automatically trace models in the worsened maps, setting the contour level as recommended by the original maps depositions. Because applying a low pass filter placed the maps beyond *DeepCryoRNA*’s range of confidently solvable resolutions this resulted in a set of 49 RNA structures (with 8 maps not yielding a usable structure), plausibly containing errors.

These low resolution structures were then compared to the corresponding high resolution (ground-truth) structures by structural alignment using a version of *Gesamt*(23) updated to handle RNA. This tool was set to handle RNA as it would proteins, but considering the C1’ atoms as representative of each base instead of the typical C*α* for each amino acid. Bases in the low resolution structures were labelled as being part of a modelling error if *Gesamt* structural alignment placed them either closest to a different base, aligned to no base at all or aligned with the correct base but at a distance (C1’ - C1’) greater than 2.75 Å. In the case where a base in the low resolution structure was placed at the location of a different ground-truth base, the distance in register between the two was also recorded and the low resolution base was flagged as potentially in a register-shift error. Bases that *Gesamt* placed within the acceptable distance of 2.75 Å from the corresponding ground-truth base were labelled as correct. This resulted in a data set of 14798 bases, of which 12089 were labelled as being correctly solved (the negative class) and 1411 were labelled as errors (the positive class), belonging to 49 low-resolution structures. Fig. 2 shows a sketch of the pipeline for creating known errors.

**Fig. 2.**
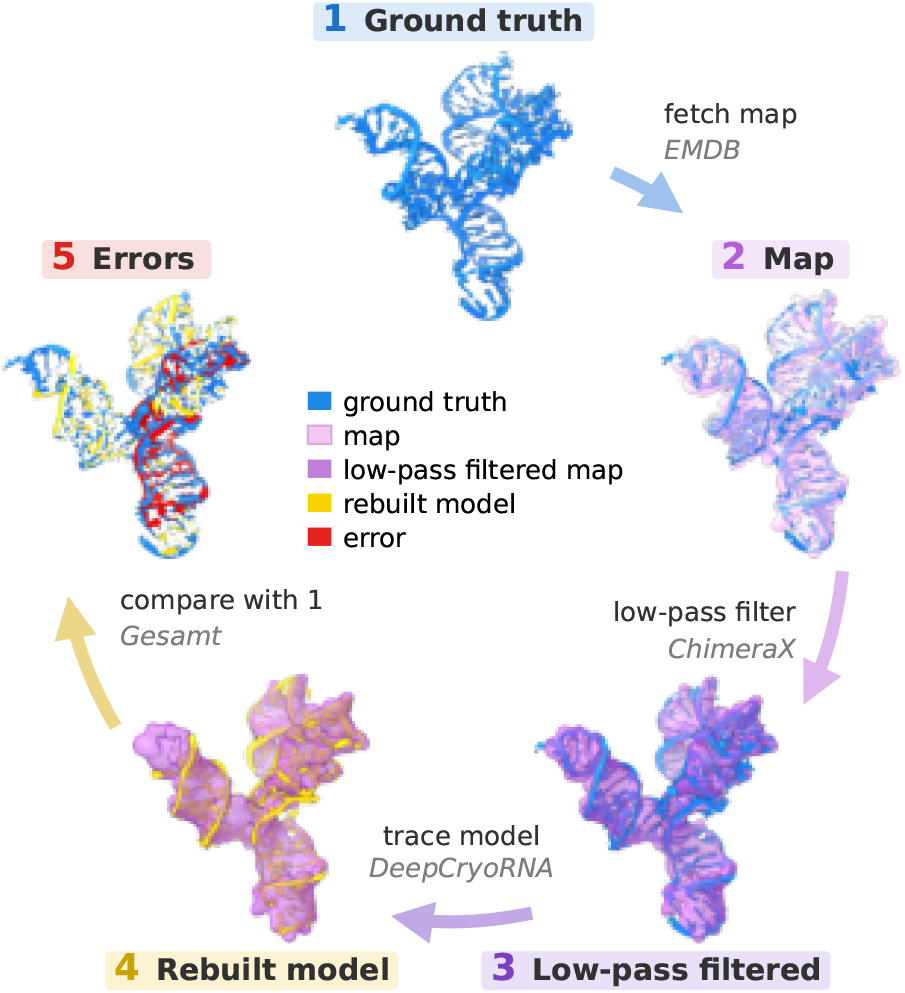
Creation of the benchmark set of known errors. (1) High-resolution single RNA chain structures determined by cryo-EM are obtained from the PDB (blue); these serve as the ground truth. (2) The corresponding Coulomb potential maps (purple) are obtained from EMDB. (3) A low pass filter is applied to these maps using *ChimeraX* (21), creating a worsened map. (4) Models are automatically built into the low-pass filtered maps using *DeepCryoRNA*, producing lower quality structures (yellow). (5) *Gesamt* is used to compare the lower quality structures to the ground truth, annotating sequence register assignment mismatches and large positional deviations as artificial errors (red).

### Prediction

For each of the 49 structures included in the error data set, the sequence was obtained from the fasta file deposited in the PDB. Using *ABCfold*(24), *AlphaFold 3* (AF3)(12), *Protenix*(14) and *Boltz-2*(25) predictions were made (using the standard *nhmmer*(26) database search), saving both the predicted coordinates and distogram in the case of AF3. Predictions were also made using *trRosettaRNA2*(18), using the provided script, which builds an MSA by a *BLASTN*(27) search of RNAcentral(28), followed by an *Infernal*(29) covariance-model search seeded with an *RNAfold*(30) secondary structure, again storing both predicted coordinates and distograms. The MSA generated for *trRosettaRNA2* was then used to also predict these targets with *RNAPro*(19). All predictions were run using an Ubuntu workstation (containing an AMD Ryzen 9 9950X × 32 CPU, a NVIDIA GeForce RTX 4080 SUPER GPU, and 192GiB of RAM).

### Contact map alignment based validation

The predicted coordinates, predicted distograms, ground-truth structures, and low resolution structures were processed into contact maps with *Conkit*(31). For coordinates (both predicted and experimentally determined), this was done using a contact definition based on the C1’-C1’ distance between two bases and a variable threshold distance *d*_*th*_. The C1’ atom was chosen to define contacts because distograms of RNA structures predicted by AF3 refer to inter-C1’ distances, and other prediction models like *trRosettaRNA2* also provide distograms based on this representative atom. For predicted distograms the conversion to contact maps was done by checking whether the centre of the bin with the highest confidence score fell below the threshold distance of *d*_*th*_, if so, a predicted contact was recorded.

As with proteins(9) *map_align* (32) was then used to calculate alignments between the contact maps of the low resolution structures and the corresponding predicted distograms, predicted structures, and ground truth structures. This tool optimises the number of contacts that overlap between the two input contact maps by introducing and extending gaps in the register(33). The resulting number of contacts is called the contact maximum overlap (CMO). In addition to calculating the CMO, *map_align* also records the necessary shifts in a contact map to achieve it, and in doing so it creates a new suggested register for the experimental structure. Bases where the register suggested in this way did not match the input register were classified as (part of) putative errors. The suggested register can be used as a guide to fix the error. Bases for which the suggested register matched the experimentally determined one were classified as modelled correctly.

To select the optimal threshold distance *d*_*th*_ for defining a contact, *conkit-validate* was used to classify all bases of the structures in the set of automatically rebuilt models, using the ground truth structure as the prediction and the re-built low resolution structure as the experiment. The threshold distance *d*_*th*_ was varied between 7.5 Å and 25 Å and putative errors were recorded. Fig. 3 shows the resulting F1 scores calculated for each distance *d*_*th*_ based on matching putative errors with known errors as true positives. The resulting F1 scores show a steep increase as *d*_*th*_ increases from 7.5 Å to ∼ 12 Å and a subsequent slow decline as *d*_*th*_ increases from ∼ 15 Å to 25 Å. Looking at more detailed statistics Fig. S1 shows that this first increase in F1 is the effect of rising recall as more necessary contacts are included while the subsequent decrease in F1 is mostly an effect of falling precision as spurious shifts become possible. Based on these F1 scores 12.5 Å was selected as the optimal threshold distance. Going forward a (predicted) contact is thus defined as a distance between C1’ atoms not exceeding 12.5 Å.

**Fig. 3.**
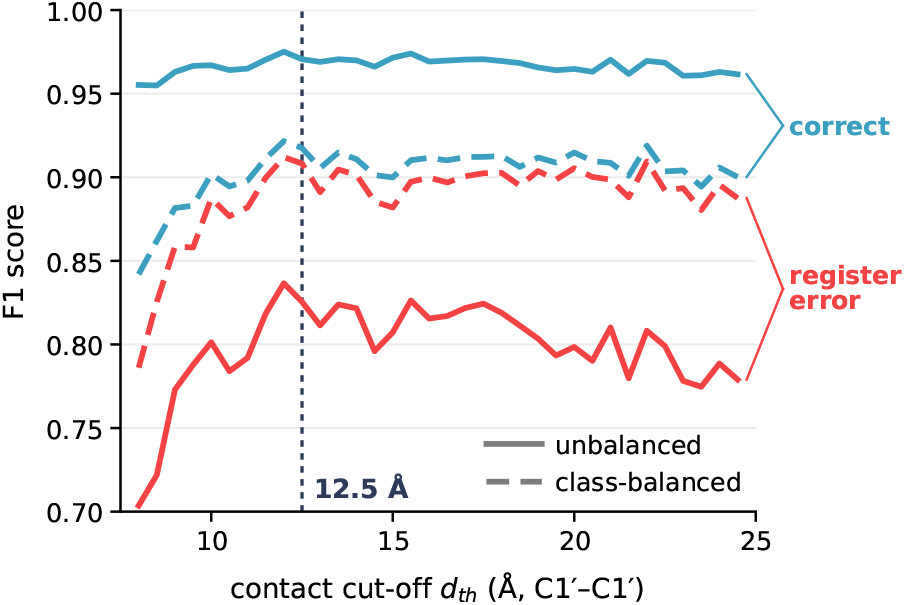
F1 score of CMO (through *map_align*) as a classifier of register errors, as a function of the contact cut-off *d*_*th*_ (C1^*′*^–C1^*′*^ distance), for the register-error and correct classes, without (solid) and with (dashed) class balancing. The dotted line marks the cut-off used, 12.5 Å.

### Machine learning classification

To also detect errors other than register shifts, machine learning classifiers were trained to detect errors using the created set of known errors.

First, a set of features were calculated to describe the similarity of the experimental and prediction based contact map locally rather than globally. These were taken to mirror *conkit-validate’s* protein-directed validation method (9). The first was the weighted root-mean-square deviation (wRMSD) defined as :

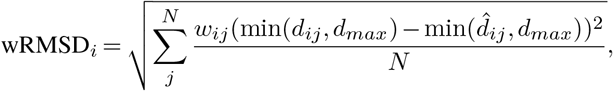

where *d*_*ij*_ is the experimental distance between residue *i* for which the wRMSD is calculated and base *j* 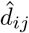, is the prediction for that same distance, *N* is the number of bases in the structure, *w*_*ij*_ is a weight based on the confidence of the prediction, and *d*_*max*_ is an upper bound on distances considered. When the input is a distogram the value of the highest bin is taken for *w*_*ij*_ and the centre of that bin for 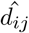, when it is a structure *w*_*ij*_ = 1 and 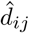 is calculated exactly from the C1’ coordinates. The final bin of a distogram is often used for overflow, gathering all predicted inter-residue distances ranging to infinity. To avoid *d*_*ij*_ becoming undefined, *d*_*max*_ was introduced and set to the lower bound of the final bin (*d*_*max*_ ≃20 Å). For similarity, when using a predicted structure we set *d*_*max*_ = 25 Å.

Next a number of features were calculated based on the correspondence between predicted and experimental contacts a base *i* makes. Here the number of contacts present in both defines *TP* , the true positives, the number of contacts made in the experiment but not the prediction defines *FP* , the false positives, the contacts made in the prediction but not the experiment defines *FN* , the false negatives, and the number of residues with which contact is made in neither defines *TN* , the true negatives. The counts FN and FP are used directly as features, and the four metrics together define the following five features:

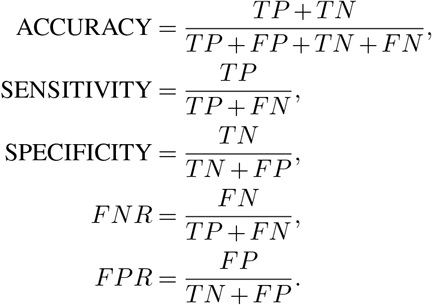

Of each of these seven features and the wRMSD versions averaged over five consecutive residues was also calculated and added to the set of features. In addition a z-score version of each feature was calculated, which compares the base to the bases within 20 Å. The z-score version of a feature *f* was defined as:

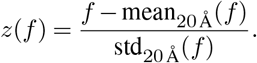

To these 24 features we then added the solvent accessibility as calculated by *areaimol*(1). Finally, to mirror the secondary structure elements used as features in the protein version of the method, we added information about the Conformational Alphabet of Nucleic Acids (CANA) classes (34) to which each residue belonged. To do this *dnatco*(35) was used to calculate the CANA class and its corresponding RMSD for each dinucleotide. 15 features were then added to each residue, one representing each CANA class and one representing no assignment. These features were set to 0 if the residue was not part of a dinucleotide assigned that CANA class, otherwise it was set to the inverse of the RMSD for that class. Because these CANA related features amounted to a lot of sparse data (features being 0), the sum of the RMSDs of dinucleotides a base belongs to was added as an additional feature.

Second, the available bases were split into a training and validation set. Prevention of data leakage from neighbouring residues was done by partitioning the entire data into blocks originating from the same structure, with consecutive residue numbers and with the same label (error or correct). For each class, error and correct, blocks were selected at random and added to the validation set until that set contained at least 20% of bases of each class, the remaining bases forming the training data set. To enable comparison between different prediction methods this was done separately for the case of prediction with AF3 structures, AF3 distograms and *RNAPro* structures, every time filtering out bases and structures which had failed to predict, or for which one of the features could not be calculated before the split. For AF3 structure prediction, this yielded a validation set of 2407 bases (consisting of 516 residues classed as errors, 20.7% of the total, and 1891 bases classed as correct, 20.3% of the total) with the remaining 9416 residues going into the training set. For AF3 distogram prediction, this yielded a validation set of 2421 bases (consisting of 499 residues classed as errors, 20.4% of the total, and 1922 bases classed as correct, 20.8% of the total) with the remaining 9278 residues going into the training set. For *RNAPro* structure prediction, this yielded a validation set of 1506 bases (consisting of 359 residues classed as errors, 20.1% of the total, and 1147 bases classed as correct, 20.1% of the total) with the remaining 5981 residues going into the training.

Third, a feature reduction scheme was applied to avoid highly correlated features. Spearman’s rank correlation coefficient *ρ* was calculated between all pairs of features across the training sets, and the features were clustered hierarchically (Ward linkage) on the distance 1− |*ρ* |, with the tree cut at a height of 0.4. From each cluster a representative feature was then taken, chosen at random. Fig. S3 shows the correlation heatmaps generated in the process. This was done separately for each prediction method (*RNAPro*, AF3 structures and AF3 distograms), and for each method once with the CANA based feature included and once without. Table S3 lists the features selected in this way.

A robust scaler was then fitted to the selected features in the training data. After which the training data were scaled and the classes were balanced by removing a number of correctly modelled bases at random. A number of machine learning classification algorithms were then trained on the balanced and scaled version of the training data using the selected features. These algorithms were: Support Vector Machine (SVM), Random Forest (RF), K-Nearest Neighbours (KNN) and LASSO linear regression classifier. The algorithms themselves were implemented through *Scikit-learn* (36). During training the hyperparameters of each method were tuned using random or grid searches depending on whether the parameter was continuous or discrete.

### Filtering

To reduce the likelihood of spuriously labelling bases as erroneously modelled, and to quantify confidence in the discovered errors, filters were developed based on three metrics not used in the error calling. First, the total number of contacts per residue is reported, to ensure our contact based methods are at all applicable. For bases which make fewer than 1 contact (excluding covalently linked neighbours) the methods are considered not useful. Second, when available the per-residue pLDDT is used as a metric of overall prediction confidence. Finally, the *Gesamt*(23) Q-score for structural alignment is calculated per block of continuous suggested register, between predicted and experimental structures. Where this score is low, putative errors might be attributed to a difference in conformation between prediction and experiment, rather than a genuine issue with the experimental structure.

In addition to calculating and reporting these metrics if possible, they were used as features to train a machine learning classifier to serve as a single filter. The filter gives each misaligned nucleotide a score; the higher the score, the more likely the call is a genuine error, and calls scoring above the filter’s threshold are reported as confident. Running average versions of the pLDDT and number of contacts were added, as well as the change in these metrics between neighbouring bases to serve as additional features, yielding seven total features for training. LASSO regression, support vector machine, and random forest algorithms were used as learning algorithms. For training, the same set of bases split off to train the machine learning classifiers using AF3 structures as the prediction was used, the set was however restricted to only the bases that were identified as misaligned using the contact-map alignment approach. The resulting set was then class balanced between True and False positives, before training. The same process was applied to the corresponding validation set to quantify the performance of the created filters, the resulting ROC-curves are shown in Fig. 4.

**Fig. 4.**
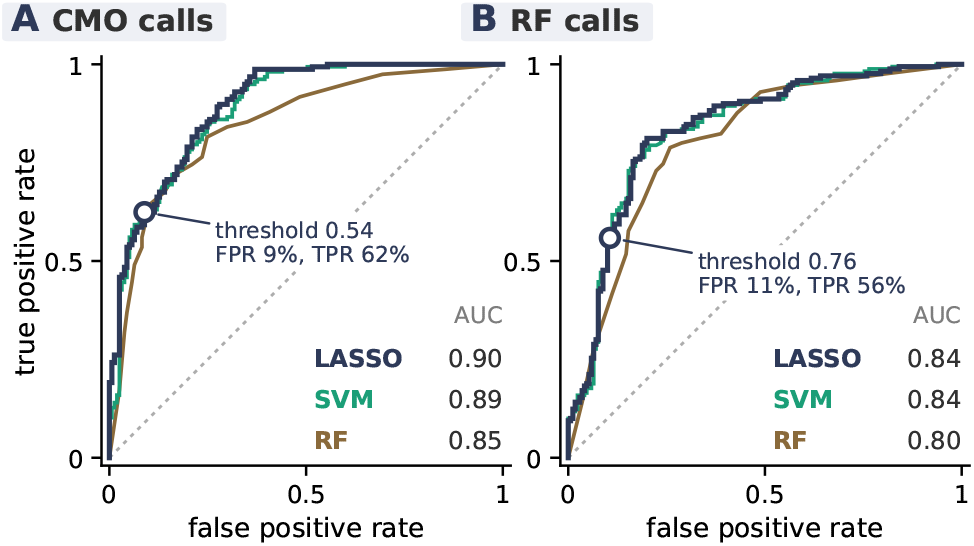
Receiver operating characteristic curves on the validation set for the falsepositive filters trained with three algorithms (LASSO, SVM and random forest), (A) for the error calls of the contact map alignment (CMO) approach and (B) for those of the random forest (RF) classifier. The dotted line represents random guessing; the area under each curve (AUC) is listed. The circles mark the thresholds set as defaults in *conkit-validate* for the LASSO filters, 0.54 (A) and 0.76 (B).

The filter created by Lasso regression shows the best performance achieving a false positive rate of 9% with a simultaneous true positive rate of 62% when the threshold is set to 0.54. This filter and threshold were set as defaults in *conkitvalidate*. A similar approach was attempted for the selected machine learning classifier (random forest), with the addition of the probability predicted by the classifier as a feature. Fig. 4 shows the ROC for the resulting filters next to those for the filters for the CMO-approach. Again the LASSO based filter performs best, albeit marginally (SVM: 0.84), now achieving 0.84 AUC compared to the 0.90 AUC found for the CMO approach. The discrepancy is likely, because the RF classifier and filter were trained on the same set, making the training data for the filter differently distributed than the validation data. The filter selected achieves a false positive rate of 11% with a simultaneous true positive rate of 56% when the threshold is set to 0.76.

For the PDB-wide scan, an additional filter was applied that acts on whole chains rather than single bases: an applicability rule for *conkit-validate* based on the *Gesamt* Q-score of the whole chain (rather than of each register block) and the average number of contacts per residue over the whole chain, this is described in Sup. Note C.

## Results

### Contact map alignment identifies known errors in the benchmark data

Fig. 5 shows how the CMO method using *map_align* performs at identifying register errors when using different prediction methods (AF3, *Boltz, Protenix, trRosettaRNA2* and *RNAPro*), where possible different input formats (structures or distogram) and the ground truth structure as a reference. Since CMO assigns no score that could be thresholded, each input gives a single operating point rather than a ROC curve; the underlying confusion matrices and the F1 score, accuracy, precision and recall are given in Fig. S2 and table S1. The results are always calculated per residue and over a set class balanced to have equal numbers of bases in a register error and correctly modelled, without applying the filter developed in Methods, Filtering. Looking at the scores obtained when using the ground truth structure provides a measure of the method’s performance in the ideal scenario: perfect prediction into the same conformation as the experiment. The method obtains an F1 score of 0.92 and an accuracy of 0.93 in this scenario. A more realistic scenario is seen when looking at the score obtained when using *RNAPro* structures, here a lower F1 score of 0.88 is achieved with accuracy falling to 0.89. Overall RNA-specific methods outscore general methods, with *RNAPro* scoring the highest.

**Fig. 5.**
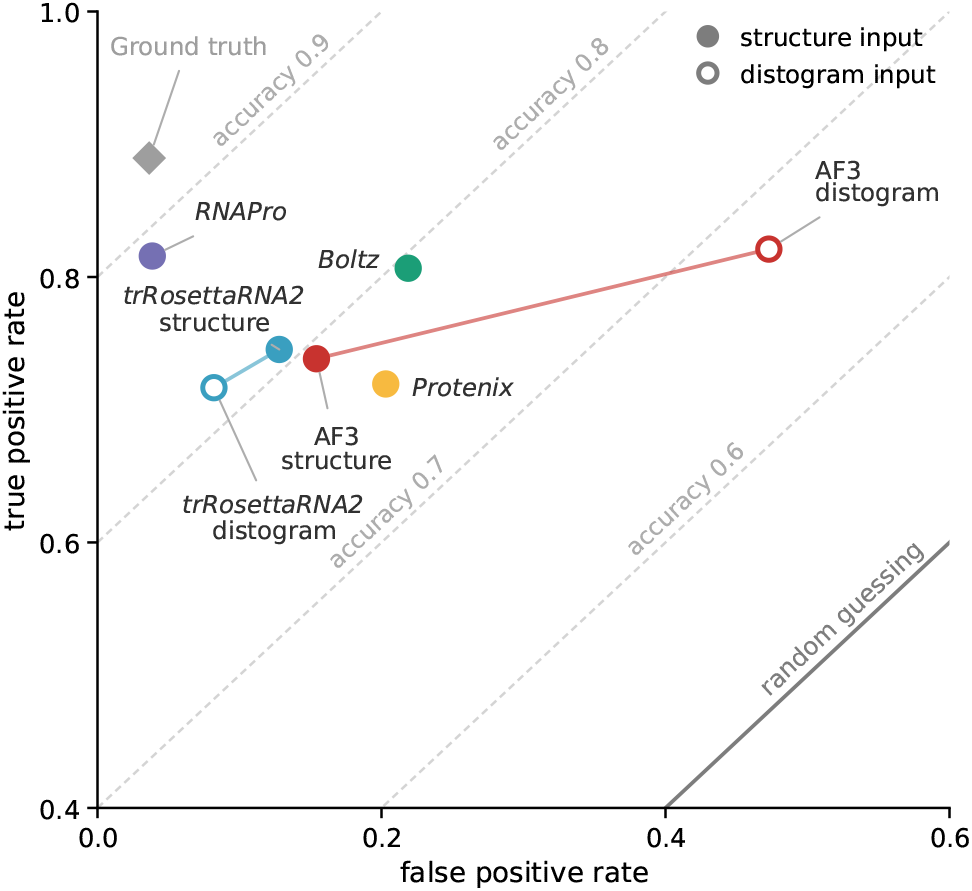
Performance of the CMO approach as a classifier of register errors for each prediction input, shown in ROC space (false positive rate against true positive rate, with register errors as the positive class). Rates are calculated per residue on a version of the benchmark data set balanced to have equal numbers of bases in register errors and correctly modelled, without the filters of Methods, Filtering. Filled markers: structure input; open markers: distogram input; lines join the two inputs of the same predictor. Colours distinguish the predictors: purple and blue for the RNA-specific models (*RNAPro* and *trRosettaRNA2*, respectively), red, yellow and green for the AF3-like models (AF3, *Protenix* and *Boltz*, respectively). Ground truth (grey diamond): the ground-truth structure used as the prediction, the best case achievable. Dashed lines: equal accuracy contours; solid line: random guessing (accuracy 0.5). The underlying confusion matrices and scores are given in Fig. S2 and table S1.

Nonetheless, using more general methods like AF3 can be preferable, due to the ability to co-fold with proteins, and because these methods are more widely available making it easier to use different scores to interpret the final result. Note that using AF3 distograms as the prediction input yields the worst results (among these methods): as Fig. 5 shows, this is a consequence of many false positive errors, with a false positive rate (0.47) more than twice that of any other input.

The new suggested register was compared to the ground truth correct register. Fig. 6 shows histograms of the difference between the suggested register and the truth among correctly identified register errors for three different prediction methods. In the best case scenario, using the ground truth structure as the prediction the suggested register was correct for 76% of residues. More realistically, when using *RNAPro* structures the register was fixed correctly 63% of the time. Like with error detection, general structure prediction methods (see table S2) performed worse than the RNA specific ones, best of this group were AF3 structures correctly fixing the register in 46% of cases.

**Fig. 6.**
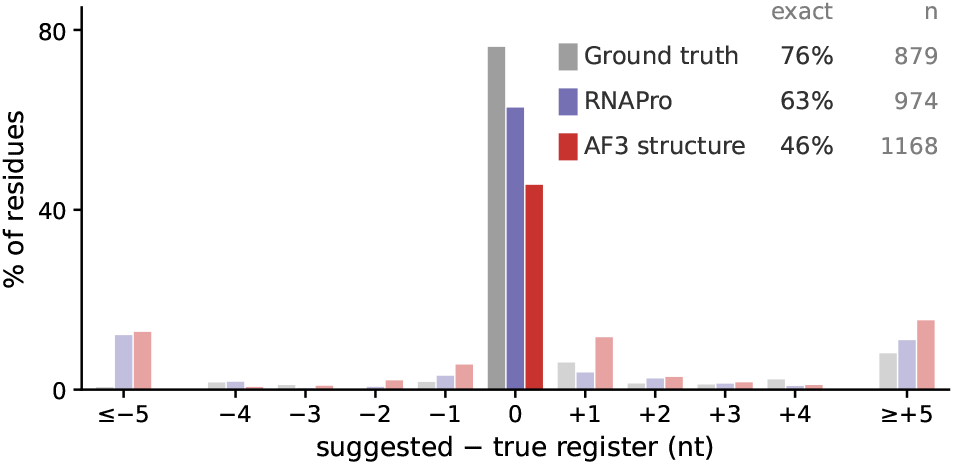
Histograms of the difference between the register suggested by the CMO approach and the ground-truth register of residues correctly classified as part of a register error. Using the ground-truth (grey), *RNAPro* (purple) and AF3 (red) structures as the input prediction. The solid central bars are the shares for which the suggested register is exactly right. Differences beyond *±*4 are pooled in the outer bars.

### Performance of machine learning classifiers

In addition to the contact map alignment approach to detect register errors, we also trained machine learning classifiers, based on four algorithms taken from *scikit-learn*(36) (RF, SVM, KNN and LASSO), using covariance-based metrics(9). This was done, separately, using features calculated with three different prediction methods and both with and without the inclusion of CANA code based features, creating a total of 24 different classifiers. To assess the relative performance of the classifiers, they were applied to a class balanced version of the appropriate validation set. The results were scored using the F1-score, the area under the receiver-operator curve (ROC-AUC), accuracy, precision and recall, and are summarised in Fig. 7 (all values in table S4).

**Fig. 7.**
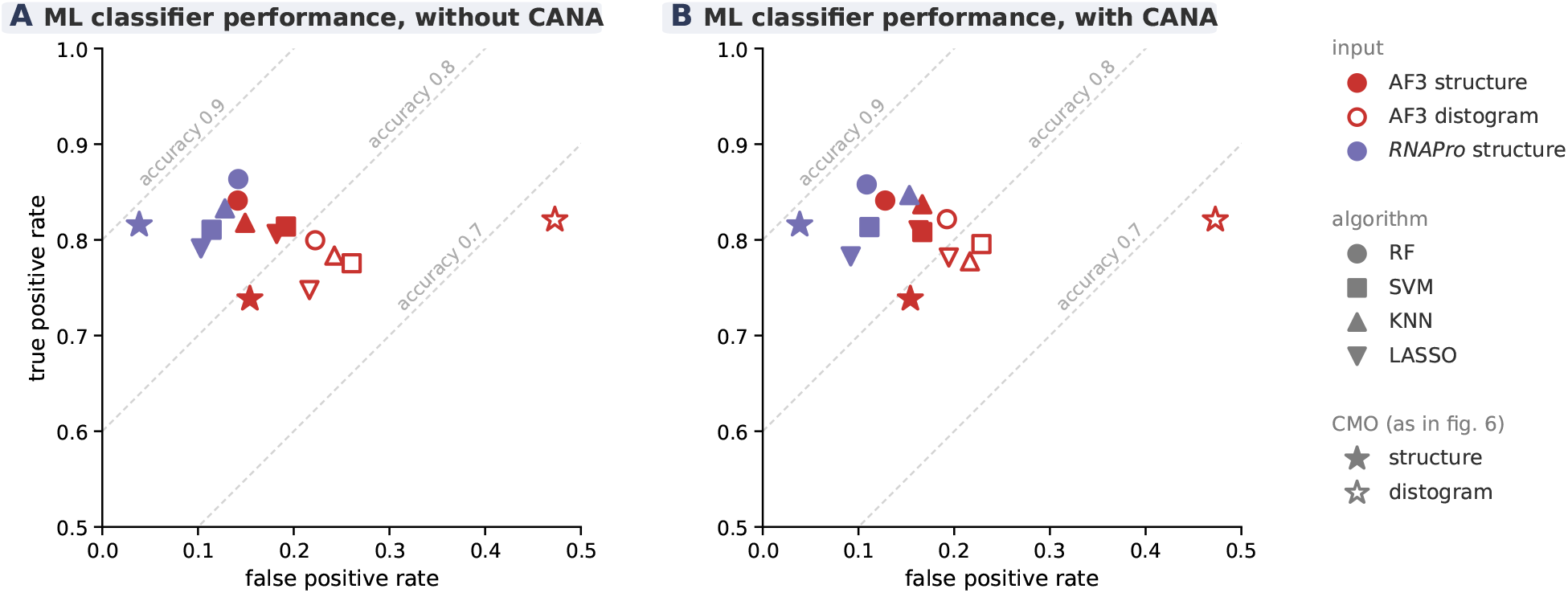
Performance of the 24 ML classifiers at their default threshold (0.5), as points in ROC space, (A) without and (B) with CANA codes, on the class-balanced validation set. Colour and fill give the prediction input as in Fig. 5 (red: AF3, purple: *RNAPro*; filled: structure, open: distogram), marker shape the learning algorithm. Stars mark the CMO approach on the same inputs (from Fig. 5); as CMO needs no training data, these were computed over the full set of artificial errors rather than the validation set only. Dashed lines are lines of equal accuracy. All metrics are listed in table S4.

The random forest classification algorithm, (RF) consistently outperformed the others across all prediction methods and both feature sets. The difference in performance when classifiers could access the CANA codes was overall smaller than the difference between algorithms and prediction methods. Among prediction methods, the structure based classifiers outperform the ones based on AF3 distograms and *RNAPro* structures slightly outperformed AF3 structures.

Because the performance improvement upon using CANA codes and *RNAPro* was only small, and considering the relative ease by which AF3 structures can be obtained (through the server(12)) we focus on discussing the results when using the RF algorithm based on AF3 structures without the use of CANA codes. Differences between this input format and others (such as distograms) are then pointed out. Fig. S4 shows the ROC for this classifier, compared to the other algorithms (and for the other inputs).

To further investigate which features are the most important to the classification, ablation studies were performed. Here the values of each feature in the validation set were randomly shuffled before reattempting prediction. For each feature the decrease in accuracy was then recorded. Per feature the process was repeated 20 times for consistency. The resulting feature importances are shown in Fig. S5 for all three prediction inputs, with and without CANA codes. For the structure-based classifiers the Sensitivity appeared to be the most important feature, across all algorithms except for KNN on AF3 structures, which relies most on the five-residue-averaged accuracy. Comparing the relative importances here to the protein case(9), most surprising is the low relative importance of the wRMSD. That feature was dominant in the SVM-based classifier trained using *AlphaFold 2* predicted distograms of proteins. The feature ranking overall second in that study: Sensitivity now appears the most useful for RNA when using structure predictions. The importance of the wRMSD reappears when RNA classifiers are trained using AF3 distograms. In that case the feature representing the correlation cluster of the sensitivity is the false positive rate, FPR, which shows little importance to the classifiers. Note that both the FPR and its smooth version were selected for these classifiers with their correlation distance slightly above the threshold in Fig. S3. The inclusion of such closely related features could cause them to appear less important to the classification task than they are.

### Complementarity of contact and map based methods in a limited PDB scan

To illustrate the value of these methods on real data, a scan of the PDB was performed. While the performance of methods presented above are independent of the experimental density maps, and thus the refinement resolution of validated structures, the set of validated structures was limited to those with a resolution better than 5 Å. This choice was made to allow a detailed comparison to CMS(20), an established map-based method. Aside from the limitation on refinement resolution, a cut-off date for deposition of 10/10/2025 was chosen, only structures determined by cryo-electron microscopy or X-ray crystallography were allowed and the set of validated entries was restricted to those that contained at least 1 RNA chain and 1 or more protein chains. The final constraint being enforced to avoid overlap with our original benchmarking data and training set. This resulted in a set of 4399 PDB entries which matched search criteria.

CMS was run on all of these structures. To perform *conkit* validation on these structures, all RNA chains were extracted, discarding any chains where the provided sequence length was below 25 (ensuring a stable fold was possible) or above 1000 bases (ensuring structure prediction was feasible) which resulted in a set of 7757 chains total. For the remaining chains structure prediction was attempted using AF3 before running the *conkit-validate* pipeline using the top predicted structure as the input. This resulted in 7580 successfully validated RNA chains, where validation on 177 chains failed largely due to crashes in the prediction model. Two of the validated chains (8UKB L7 and L8, neither flagged by any method) were removed from the PDB as superseded by entry 9PKB between the original construction of the data set and final computations for this work, so figures 8 and 9 cover the remaining 7578 chains.

**Fig. 8.**
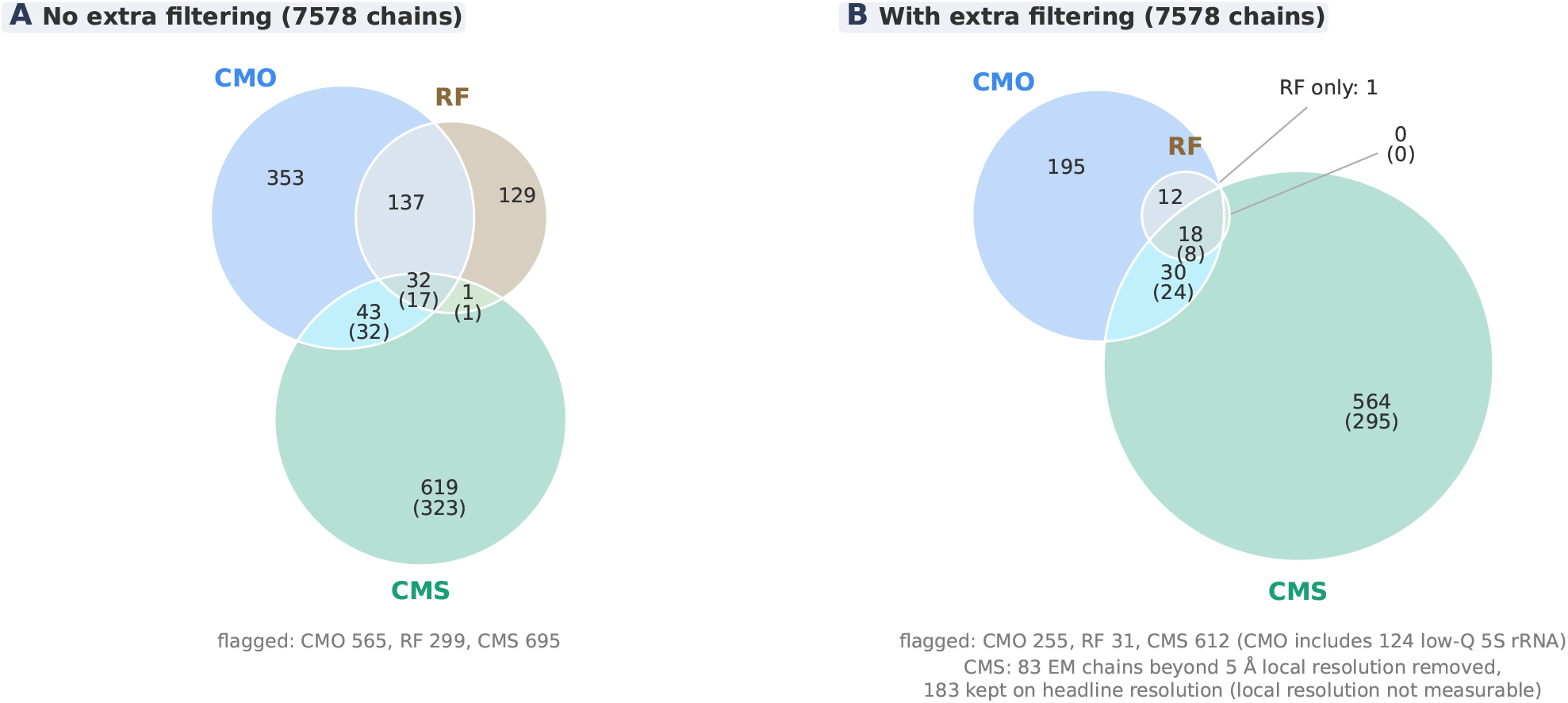
Chains flagged as containing putative errors by *conkit-validate* using the CMO (blue) and the random forest classifier (RF, brown), and by *checkMySequence* (CMS, green, any error type), out of the 7578 chains validated by *conkit-validate*. A: without additional filtering. B: with the additional filters described in Supplementary Section C: CMO and RF flags are only counted for chains passing the applicability rule (global Q-score *≥* 0.30 and *≥* 1.45 contacts per residue), CMS flags only for chains within its 5 Å working range, judged for EM chains by the local resolution of the chain itself (83 chains removed); 183 flagged EM chains whose local resolution could not be measured are kept on their headline resolution. Of the 255 filtered CMO chains, 124 are low-Q 5S rRNA chains (Supplementary Section C). The bracketed number in every region CMS appears in gives how many of its chains were flagged specifically for a register shift. A version leaving out EM chains without a measurable local resolution is shown in Fig. S7.

**Fig. 9.**
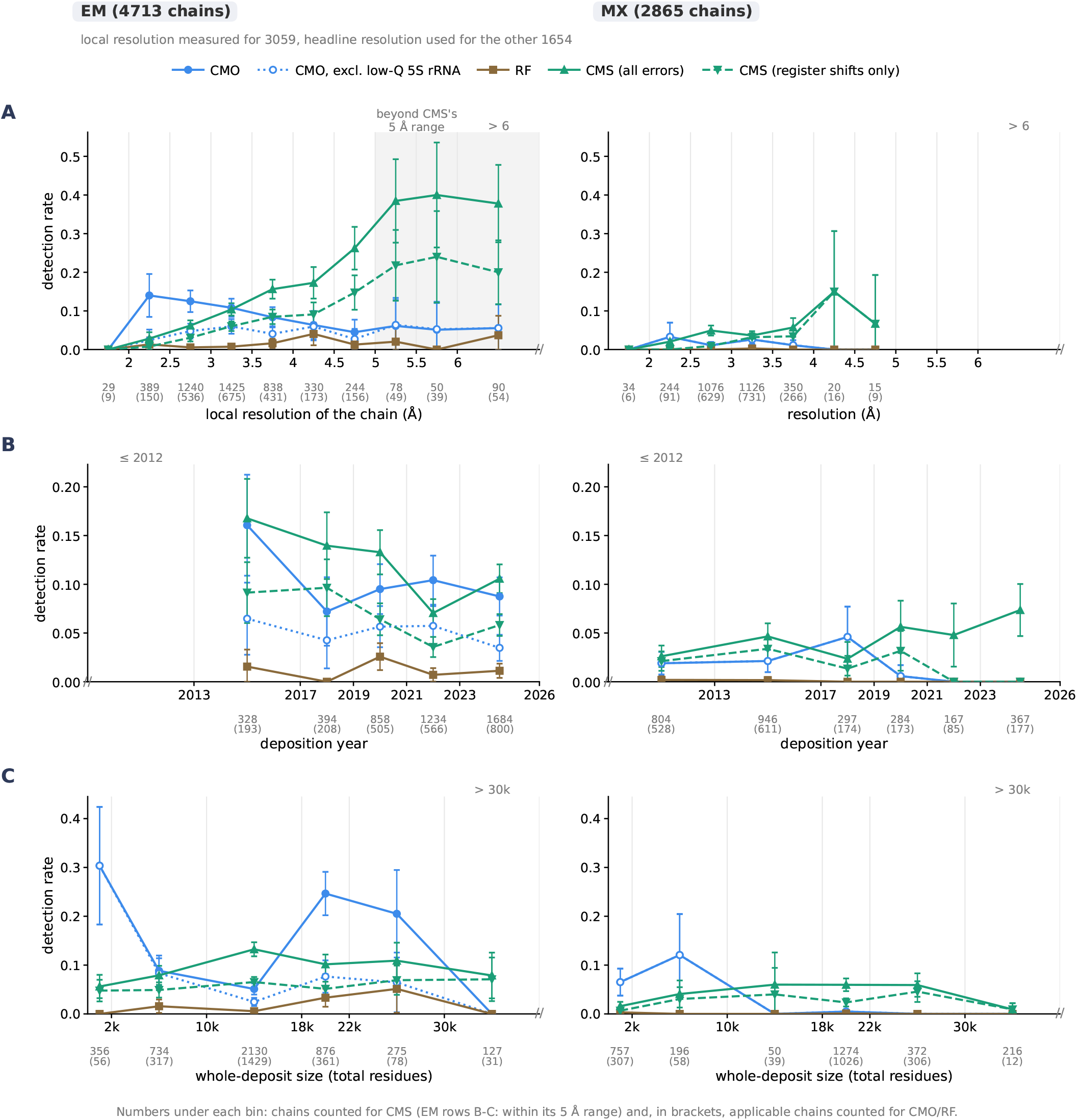
Detection rate of putative errors by CMO (blue, solid; dotted: excluding low-Q 5S rRNA chains), RF (brown), CMS (green) and CMS considering only register-shift errors (green, dashed), for structures determined by electron microscopy (EM, left) and X-ray crystallography (MX, right), against (A) resolution, (B) deposition year and (C) whole-deposit size (total number of modelled residues in the entry). All panels use the additional filtering of Fig. 8: CMO and RF rates are computed over applicable chains only. For EM, resolution is the local resolution of each chain’s own density where it could be measured (3059 of 4713 chains) and the headline resolution otherwise; in (A) bins beyond CMS’s 5 Å working range are shaded, in (B) and (C) CMS only counts chains within that range. Numbers under each bin: chains counted for CMS and, in brackets, applicable chains counted for CMO/RF; error bars: 95% intervals. A version leaving out EM chains without a measurable local resolution is shown in Fig. S8.

After applying the trained false positive filters and imposing a minimum error size of 6 consecutive bases, 565 chains were flagged as containing putative errors by the CMO method while the Random forest classifier flagged 299 chains. The map-based CMS identified 695 chains in that set as containing putative errors of any type, of which 373 were specifically register-shift errors, the error type *conkit-validate* targets. Fig. 8A shows how the sets of chains containing putative errors identified by different methods relate to each other, with the register-shift-only subset of CMS’s calls broken out within each region it appears in.

Visual inspection of a number of the chains in which *conkit-validate* flagged an error, revealed large regions of these chains to be unravelled, or to have globally different conformations able to trip up CMO and RF validation. Despite the application of the local residue level filter developed above some of those calls are likely false positives. To remedy this the *Gesamt* Q-Score aligning each chain with its AF3 prediction was calculated along with the average number of contacts per residue. Using an active learning protocol detailed in Supplementary Section C, chain level thresholds (*Q*≥ 0.3 and contacts per residue ≥1.45) were set on these two properties to determine whether the contact based methods are applicable. Of the 7556 chains for which both could be computed, 4020 pass this rule. The 22 chains for which values could not be computed are treated as not applicable.

Likewise visual inspection of the maps corresponding to chains in which errors were identified by CMS, showed that in the case of cryo-EM the region of map into which those chains were built had visually poorer resolution than the rest of the structure. Because CMS is only reliable at resolutions better than 5 Å and the local resolution for some of these chains might fall below that threshold, local resolutions were calculated per chain for all validated cryo-EM structures that had half-maps available. The protocol for this calculation is given in Supplementary Section C.

The effect, on the amount of confidently found errors, of applying this extra applicability filtering, based on global conformation and number of contacts in the case of CMO and RF and local rather than global resolution in the case of CMS is shown in Fig. 8 B. Note that in the case of CMO and RF, in addition to this filtering, a set of 5S rRNA chains was identified which while having a globally similar structure to the AF3 prediction, differed enough in a single stem-loop to cause false positives to show up there. With the additional filtering, CMO keeps 255 of its 565 flagged chains and RF 31 of its 299, while 83 of the chains flagged by CMS lie beyond its working range, leaving 612. Of the 255 remaining CMO calls 124 fall in the set of 5S rRNA chains. Because these are identified as clear false positives, but cannot be excluded with either of the developed filters, numbers both with and without this set included are shown in both Fig. 8 and in Fig. 9. Adding this filtering step also removed all but one call made by the RF classifier not corroborated by CMO. This is consistent with the finding that for protein most ML classifier (SVM) calls not supported by CMO were cases of global conformational mismatch between the prediction and deposit.

To identify trends in the error rate, the results of the PDB scan are shown plotted against (local) resolution, deposition date and total size of the deposit in Fig. 9. Every method flags a larger share of EM chains than of MX chains, and in MX structures the rates of putative errors stay low throughout. For both MX and EM structures, the rate of putative errors identified by CMS rises as the resolution becomes worse, with the sharpest rise occurring near the 5 Å cutoff. After that cutoff, for EM structures the rate of errors levels to a plateau of about 40%, where the method is no longer reliable. No MX structures were validated with a resolution worse than the 5 Å cutoff. The CMO method shows an initial rise in putative errors among EM structures as the resolution worsens, with the rate detected (when excluding the set of unreliable 5S rRNAs) closely tracking the rate of errors detected by CMS before levelling out around 3 Å with ∼5% of chains having putative errors at worse resolutions. The CMO error rate in MX structures shows no trend, hovering between 0% and 3%.

Over time, putative errors in EM and MX structures show opposing trends as shown in Fig. 9B. CMS finds fewer errors in newer EM deposits, potentially due to increasing resolution, while the CMO rate excluding the low-Q 5S rRNA chains stays between 4% and 6% throughout. In MX structures, CMS shows a growing rate of total errors while the number of errors found by CMO and the specific register shift errors identified by CMS have collapsed since 2021.

The clearest feature when looking at the correspondence between error rate and the total number of modelled residues (in the deposit, not just the flagged chain), is an increase in CMO identified putative errors for deposits with between 18k and 30k residues. However, this bump in error detection rate disappears when excluding the set of 5S rRNAs previously flagged as likely false positives. When the difficulty in predicting the structures of these specific 5S rRNAs is taken into account, Fig. 9C shows the only correlation between deposit size and error rate is an increase in CMO calls for smaller deposits.

Below case studies are provided to illustrate some confirmed true positives but also one false positive and a case where no other method could confirm or refute the error.

### Case study from the PDB 4WCE chain Y: a known error that map-based cross-validation could not confirm

To illustrate the potential of *conkit-validate* to identify registershift errors in RNA, some cases from the PDB scan were selected. For these cases we looked for independent evidence of the putative error. The first selected case was chain Y of PDB entry 4WCE(37), a 5S rRNA (50S subunit) in the *S. aureus* ribosome. This structure was solved through X-ray crystallography at a resolution of 3.5 Å. In this structure *conkit-validate*’s CMO method flagged a window between bases 80 and 113 as having a −1 register shift error (Fig. 10B). The RF classifier also scores this region as having a higher than average likelihood of containing erroneous bases, with the score averaging to 0.49 across the window (ranging from 0.17 to 0.75). While the precise signal of the classifier method remains noisy and the corresponding filter never clears its 0.76 threshold anywhere in the window (mean 0.07, range −0.54 to 0.51), the filter for the CMO method indicates this is a likely true positive, remaining comfortably above its 0.54 threshold throughout (mean 0.94, range 0.61 to 1.05). Note the filter scores are reported by *conkit-validate* in the command line but not plotted in Fig. 10in it’s figure output, instead hashing indicates if a region flagged as a putative error has a confidence lower than the preset threshold. Fig. 10 shows *conkit-validate*’s output.

**Fig. 10.**
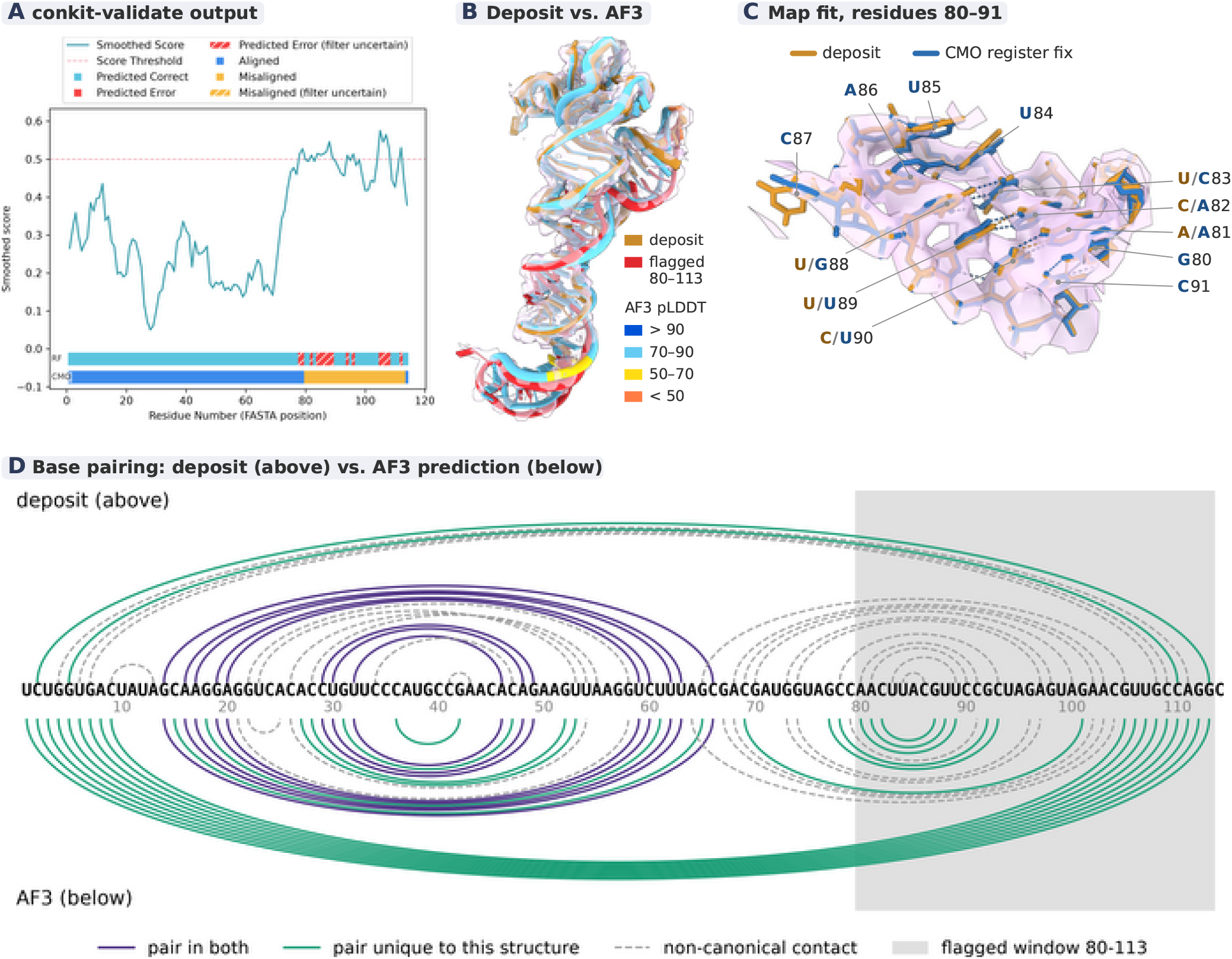
Case study figure for 4WCE chain Y. A: *conkit-validate*’s output plot (cyan line: smoothed RF score, dashed red line: threshold for the RF score, top (red/cyan) track: RF error call, and bottom (blue/yellow) track: CMO error calls) for chain Y, hatched segments are calls the false-positive filter rates as uncertain (below its threshold). B: the deposited chain (ochre, with the flagged register-shift window 80-113 in red) superposed with the AF3 prediction rigid-body fitted onto the deposit via the stable anchor region 1-79, coloured by per-residue pLDDT (key next to the structure), in the omit map (pale lilac surface, shown within 2.6 Å of either model). C: close-up of the loop residues 80-91 of the deposit (ochre) and *conkit* ‘s CMO call (hypothesis 2, after refinement; blue) in the omit map ; hydrogen bonds in each model are shown as dashed lines. D: arc diagram of the base pairing in the deposit (above the sequence) and the unrefined AF3 prediction (below); purple arcs are pairs present in both structures, green arcs pairs present only in that structure, and dashed grey arcs implausible non-canonical contacts The grey band marks the flagged window.

This case turns out to have been flagged previously in the literature (4). In that study the map-based validation software *DoubleHelix* (now part of CMS) flagged a potential register shift of 1 base starting from base 81. Just like the *conkit* result, this was found consistent with the deletion of one nucleotide in a terminal hairpin loop. The underlying cause of the error is the sequence: the deposit uses the 114-nucleotide sequence which is a minority copy of the relevant 5S rRNA gene instead of the 115-nucleotide majority copy. The modelled sequence thus lacks one nucleotide near position 81, relative to the map. This pulled all following residues into a −1 register shift.

A comparison between the base pairing of the deposit and the AF3 predicted structure, Fig. 10 D, shows AF3’s register results in more canonical (Watson-Crick or wobble) pairs. Across the hairpin stem and closing helix (the affected region, covering nucleotides 1 to 9 and 77 to 113) AF3 forms 14 canonical pairs, while the deposit only makes 2.

Density-based cross-validation of the error, and of *conkit*’s suggested fix, was attempted to confirm the veracity of the error. Four hypotheses were put forward for the correct structure of 4WCE chain Y. First, the exact deposited structure was proposed to serve as a baseline comparison. Second, *Coot*’s(38) `mutate_base` feature was used to change the base identities exactly according to *conkit*’s CMO call, keeping the exact backbone from the deposit (in this window the shifted minority sequence matches the majority copy). Third, the structure (for the minority copy) predicted by AF3 was placed into the map using a rigid-body fit to the region of the chain not flagged by *conkit*.Fourth, AF3 was used to predict the structure of the majority copy, the resulting model was placed using the same method. An omit map was constructed to avoid X-ray model-phasing bias in favour of the deposited structure: the residues of the flagged window were deleted from the deposited model and *σ*_*A*_-weighted map coefficients were recomputed from the remaining coordinates with *REFMAC5*, without further refinement, so that the map over the window carries no contribution from their deposited positions. Subsequently all four proposed structures were refined into that map using *coot-mini-rsr*, applying a staged weight-ramp protocol. The real-space correlation coefficient (RSCC) was then computed over the flagged window as a whole at the end point of each of the four refinement runs, and the difference in RSCC (ΔRSCC) between the raw, unrefined deposited structure and the refined end points of all four hypotheses (including the unmutated baseline itself, hypothesis 1) was used to test the relative suitability of each proposed structure. Resulting ΔRSCC numbers are shown in Table 1.

**Table 1.** Change in real-space correlation coefficient over the flagged window of 4WCE chain Y (residues 80–113, computed as one region), relative to the raw, unrefined deposited structure (RSCC 0.650), for the deposited baseline and three proposed corrections. Hypothesis 1’s change is the effect of the refinement pipeline alone (refinement with no hypothesis applied) and serves as the baseline the other rows are judged against. Hypothesis 2 does not differ meaningfully from it, hypothesis 3 drops about four times as much, and hypothesis 4, which repeats hypothesis 3 with AF3’s prediction for the majority gene copy, falls in between.

| hypothesis | ΔRSCC |
|---|---|
| 1: deposited baseline | −0.028 |
| 2: *conkit*’s CMO call (mutate_base) | −0.019 |
| 3: AF3 rigid-body fit (deposited sequence) | −0.119 |
| 4: AF3 rigid-body fit (majority sequence) | −0.071 |

Of the proposed alternative structures, the prediction of the minority copy fits the map worst. The prediction based on the corrected sequence, the majority copy does better, showing the map prefers a prediction of the majority copy over the minority copy, but still this hypothesis scores below the baseline. The fix that changed the base identity using `mutate_base` but kept the backbone tracing performs best, doing better than the deposited structure when run through the *coot* refinement, but not when compared to the exact deposit before additional refinement. Based on map fit alone, the baseline and the fix suggested by the second hypothesis can thus not be distinguished (Fig. 10C).

Further investigation of the literature on this case reveals CMS(20) flags errors in two nearby protein chains. An improved fit was only reported once all three errors were fixed simultaneously(4). Because the simultaneous detection and fixing of errors in multiple chains are currently beyond the scope of *conkit-validate*, we refrain from redoing the calculation. Instead, we note this presents a limitation to *conkit*’s error-fixing method: even a clear true positive error can be hard to verify on a model-map-fit basis when neighbouring chains affect the fit. Crucially, the error in this case was propagated to 13 later deposits. This shows the importance of fixing such errors at source to avoid introduction of errors by methods which use existing structures (or homologs) as a starting point.

### Case study from the PDB 8RXH chain S2: register-shift error confirmed by both density and independent cross-structural precedent

The second selected case was that of chain S2 of PDB entry 8RXH(39), a structure associated with a ribosome of *Leishmania major*, solved by cryo-electron microscopy at a reported resolution of 2.93 Å. Here, *conkit* flagged a putative +1 register error between residues 47 and 53 using CMO (Fig. 11B). The RF method again shows a more complicated signal, but does flag several residues in the same window as being erroneously placed, as shown in Fig. 11A. CMS also identifies this chain as likely containing a register shift error, although it puts forward a far larger −6 shift in a far larger window (between 37 and 66). A corrected model was then built by inserting a nucleotide after U47 and removing U55. Correspondingly the sequence identity of the residues in between was adjusted to the correct register, which changed the base identities of the nucleotides at positions 49 (C becoming U), 51 (U becoming G) and 53 (G becoming U). *Coot*’s `mutate_base` was then used to construct or complete the altered and inserted nucleotides. The resulting model was then refined into the map using *ISOLDE*(40) in *ChimeraX*(21). This was done, first while applying base pairing restraints based on the pairing from the AF3 structure while the map weight was ramped up, then the restraints were turned off and the map weight was ramped up again from a low weight. Finally, the refinement process left the base of C48 lying flipped out outside the density due to the rebuilding process, while the density occupied by C48 in the deposit was left empty. This base was moved into the density using temporary position restraints and remained there after the restraints were released. This resulted in a rise in RSCC across the whole relevant window (residues 46 to 57), relative to the unrefined deposit, of 0.043 (rising from 0.242 for the deposit to 0.285 for the suggested fix). In the corrected region this RSCC gain was approximately twice as large as the gain of the chain as a whole (0.024), which suggests this is an effect of the fixed register rather than an artefact from the refinement pipeline alone. Fig. 11 C shows the fixed structure of the corrected region overlaid on the deposited one.

**Fig. 11.**
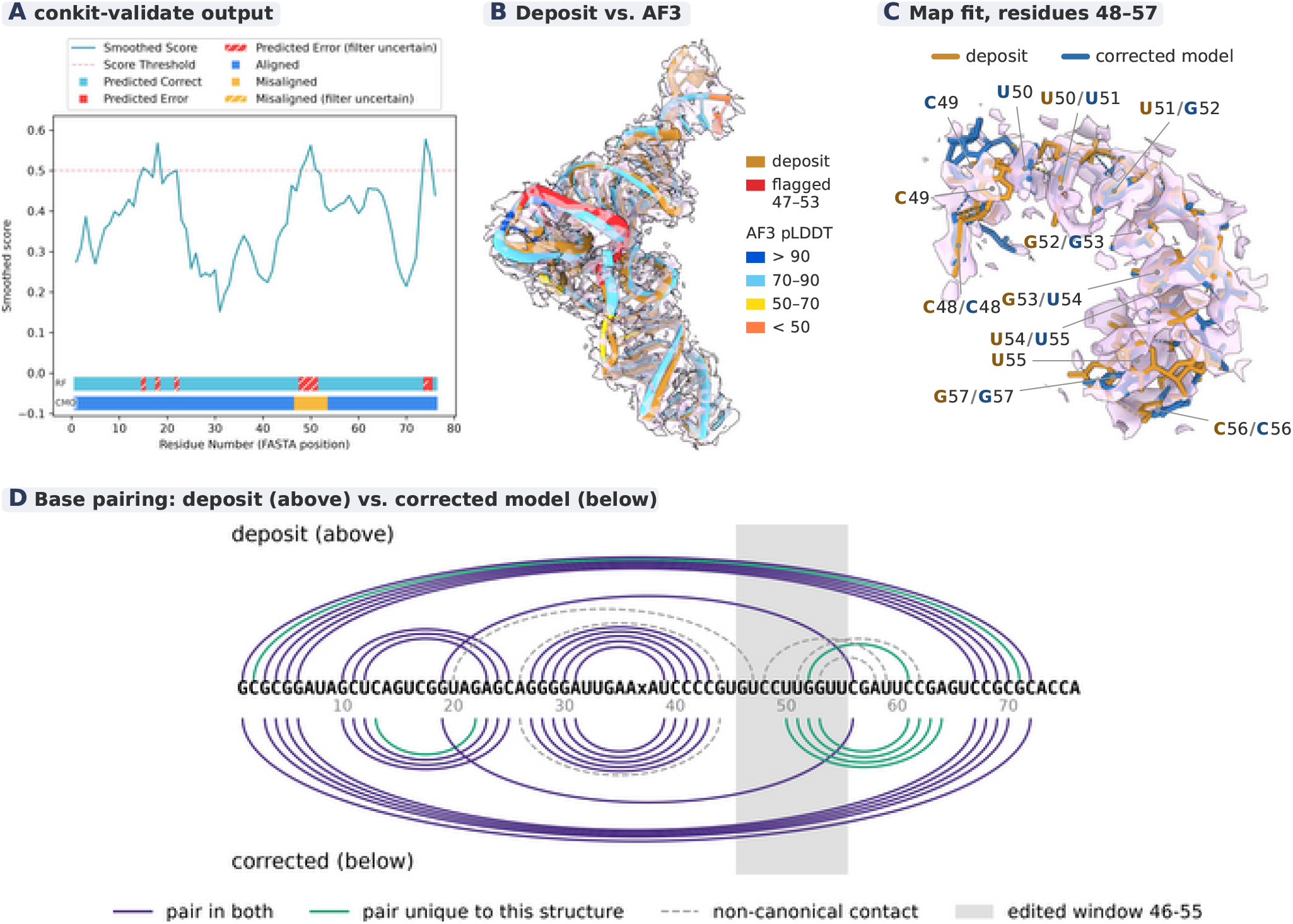
Case study figure for 8RXH chain S2. A: *conkit-validate*’s output plot (smoothed score, RF and CMO tracks) for chain S2. B: the deposited chain (ochre, with the putative register-shift window 47-53 in red) superposed with the AF3 prediction (coloured by per-residue pLDDT) in the map (pale lilac surface, shown within 2.6 Å of either model). C: close-up of residues 48-57 of the deposit (ochre) and the corrected model (blue) in the density map (shown within 2.6 Å of either model)D: arc diagram of the base pairing in the deposit (above the sequence) and the corrected model (below), in which the register was shifted by inserting a nucleotide after U47 and removing one at position 55, leaving the sequence unchanged, and which was refined into the map (see text). purple arcs are pairs present in both structures, green arcs pairs present only in that structure, and dashed grey arcs implausible non-canonical contacts The grey band marks the edited window (46-55).

Two further lines of evidence support the correction: First, base-pairing analysis shows the corrected register converts implausible non-canonical contacts into canonical pairs in the affected window (Fig. 11D), with the correction forming four canonical pairs while the deposit only had one in this region. Second, a sequence search of the PDB revealed 18 other PDB entities with the exact same sequence. None of those other entities shares the register assignment of 8RXH S2, while they do all agree with each other and the AF3 prediction.

Notably, this has the reverse implications from the previous case study: the error did not contaminate the rest of the PDB.

### Case study from the PDB 7L48 chain E: an inconclusive result that illustrates the limits of the density-based check

A third selected case was chain E of PDB entry 7L48(41). Unlike the previous two cases, this chain was not part of a ribosome-scale assembly. Rather, the entry was a complex between one protein, Cas12f, and an sgRNA. The structure was determined by cryo-EM at a resolution of 3.9 Å. *Conkit*’s CMO flagged two windows as having a +2 register shift, from base 133 to 140 and again from base 182 to 190. The RF classifier also flagged these residues, scoring an average of 0.909 across window 1 and 0.900 across window 2 (both above the threshold), as shown in Fig. 12.

**Fig. 12.**
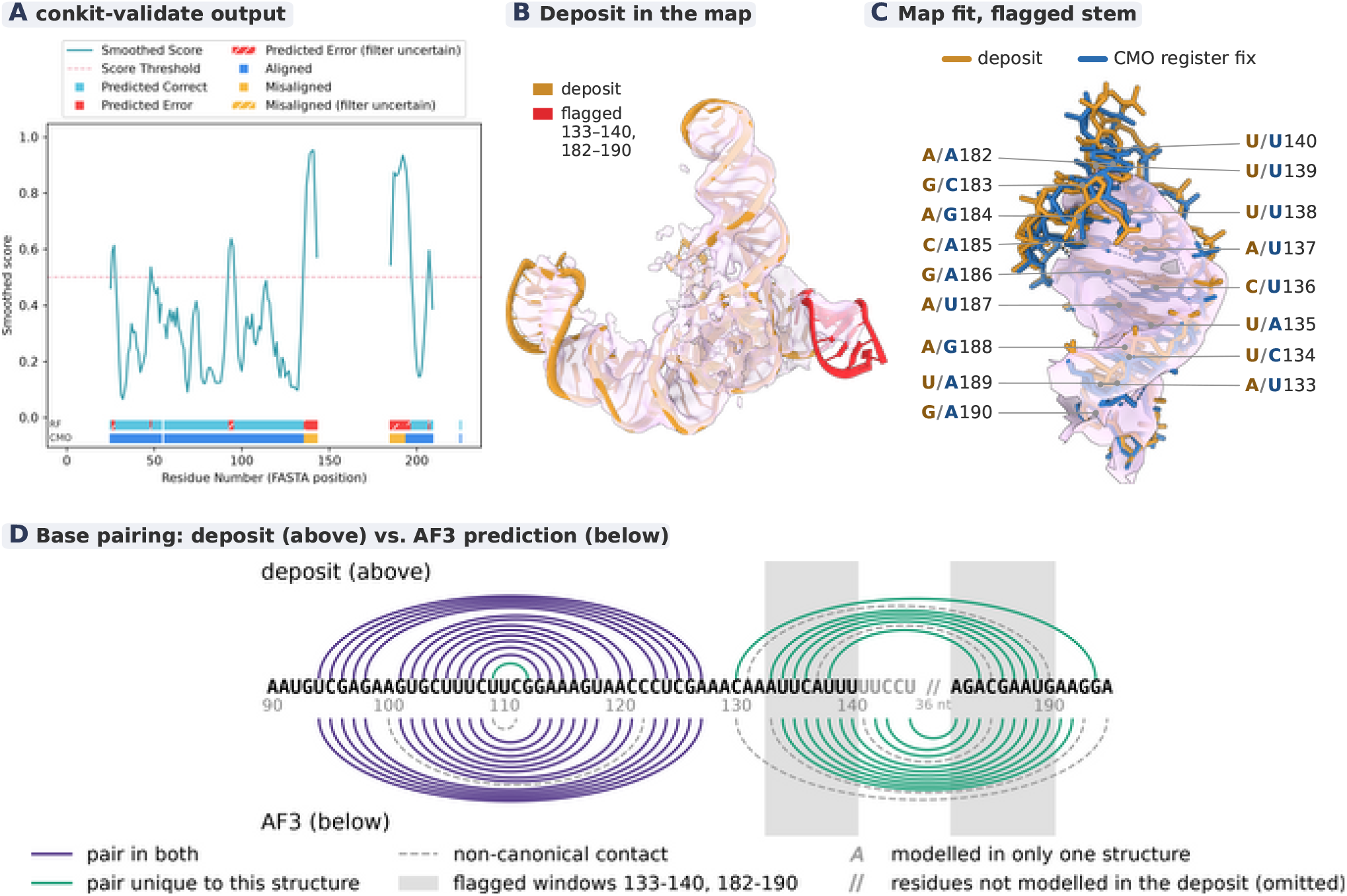
Case study figure for 7L48 chain E. A: *conkit-validate*’s output plot (smoothed score, RF and CMO tracks) for chain E. B: the deposited chain (ochre, with the two flagged windows 133-140 and 182-190 in red) in the density map (pale lilac surface, and shown within 6 Å of the chain)The rest of the sgRNA sits inside a continuous envelope, while the stem formed by the two flagged windows lies largely outside it. C: close-up of that stem in the deposit (ochre) and with *conkit* ‘s CMO call applied (the +2 shift in both windows, hypothesis 2 in Table 2, after refinement in blue), in the same map (shown within 2.6 Å of either model)Both registers fit the weak density about equally. D: arc diagram of the base pairing in the deposit (above the sequence) and the unrefined AF3 prediction (below), drawn as in Fig. 10, for residues 90-195. Residues the deposit does not model are omitted (marked //), except 141-145, which AF3 pairs with the second flagged window (grey letters). The grey bands mark the two flagged windows.

Map-based validation with CMS, however, did not identify the chain as containing a putative error. Q-scores(42) (calculated through *ChimeraX* `qscore`(21), Supplementary Section D) show the windows are modelled into poorer density than the rest of the chain (median Q-scores of 0.34 and 0.33 against 0.48 for the rest of the chain).This weak density might explain why CMS fails to flag the region.

**Table 2.** Change in real-space correlation coefficient, relative to the raw, unrefined deposited structure, for the deposited baseline and four proposed corrections to 7L48 chain E, computed over window 1 (residues 133–140), window 2 (residues 182–190) and both windows together, each as one region (raw deposit: 0.447, 0.426 and 0.568). As in Table 1, the deposited baseline’s own change (row 1) is the effect of the refinement pipeline alone and serves as the baseline the other rows are judged against. None of hypotheses 2–5 differs from it by more than 0.04 in any column, and none improves on it consistently.

| hypothesis | window 1 | window 2 | total |
|---|---|---|---|
| 1: deposited baseline | +0.058 | −0.031 | −0.001 |
| 2: +2 shift, both windows | +0.044 | −0.014 | +0.006 |
| 3: +4 shift, window 1 only | +0.016 | −0.028 | −0.012 |
| 4: +4 shift, window 2 only | +0.033 | −0.047 | −0.003 |
| 5: +4 shift, both windows | +0.044 | −0.044 | +0.013 |

Visual inspection of the structure revealed that the two windows are not separate errors; in fact, they form a base-paired stem together. Additionally, the bases between 140 and 182 (between the windows) were left unmodelled in the structure, due to missing density. Due to these two facts, and the consistency of the shift between them, the two windows were treated as a single error. The edges of this single error can be described as a deletion of two nucleotides before the start of the first window (occurring near position 132), followed by an insertion of two nucleotides near the close of the second window (near position 191). All nucleotides in between, including both windows and the unmodelled loop between them, end up shifted by +2.

The two windows forming a paired double helix brings the additional wrinkle that, on a pure contact map basis, a +2 shift in both windows would look indistinguishable from a +4 shift in either window alone. Base pairing (Fig. 12D) shows that indeed a total shift of +4 of pairing partners improves the number of canonical pairs, with the AF3 prediction showing 10 canonical pairs where the deposit only managed 5. For this reason, five different hypotheses were put forward. Like before, first the null-hypothesis of the deposited register, second *conkit*’s strict +2 register shift across both windows, third a +4 shift applied only to the first window, fourth a +4 shift in only the second window, and last a +4 shift in both windows, which is not strictly compatible with *conkit*’s suggested register. Constructing each of these (except the null-hypothesis) using `mutate_base` and then refining as above with *coot-mini-rsr* produced the ΔRSCC shown in Table 2.

None of the alternative hypotheses appears to meaningfully improve the model-map fit compared to the deposit (Fig. 12C). Neither do they worsen the RSCC, by more than the baseline change. This implies that, as map-based validation did not reject the deposit either, the map cannot distinguish the different registers in this case. Unlike for 8RXH, the PDB contains only 7L49 from the same study(41) modelling the flagged region of this sgRNA, which has the same register as 7L48. Register information for such cases thus needs to be informed by mapless methods (like *conkit-validate*). While the veracity of this error remains unconfirmed, this case highlights the need for good mapless validation methods, especially in structures or regions of structures built into poorer density.

### Case study from the PDB 8I3Z chain B: a false positive resolved by adding structural context

The fourth selected case was that of chain B in PDB entry 8I3Z(43), an NAD-II riboswitch bound to NMN solved at a resolution of 1.67 Å by X-ray crystallography. *Conkit* flagged a window from base 30 to base 48 as having a +5 register shift using the CMO method. The RF classifier scored these same bases at an average of 0.87, well above its threshold of 0.5 (higher scores indicate a more likely error), as seen in Fig. 13. As before, *conkit* was run providing an AF3 prediction of just chain B, not chain A from the same deposit or the ligand present in the structure.

**Fig. 13.**
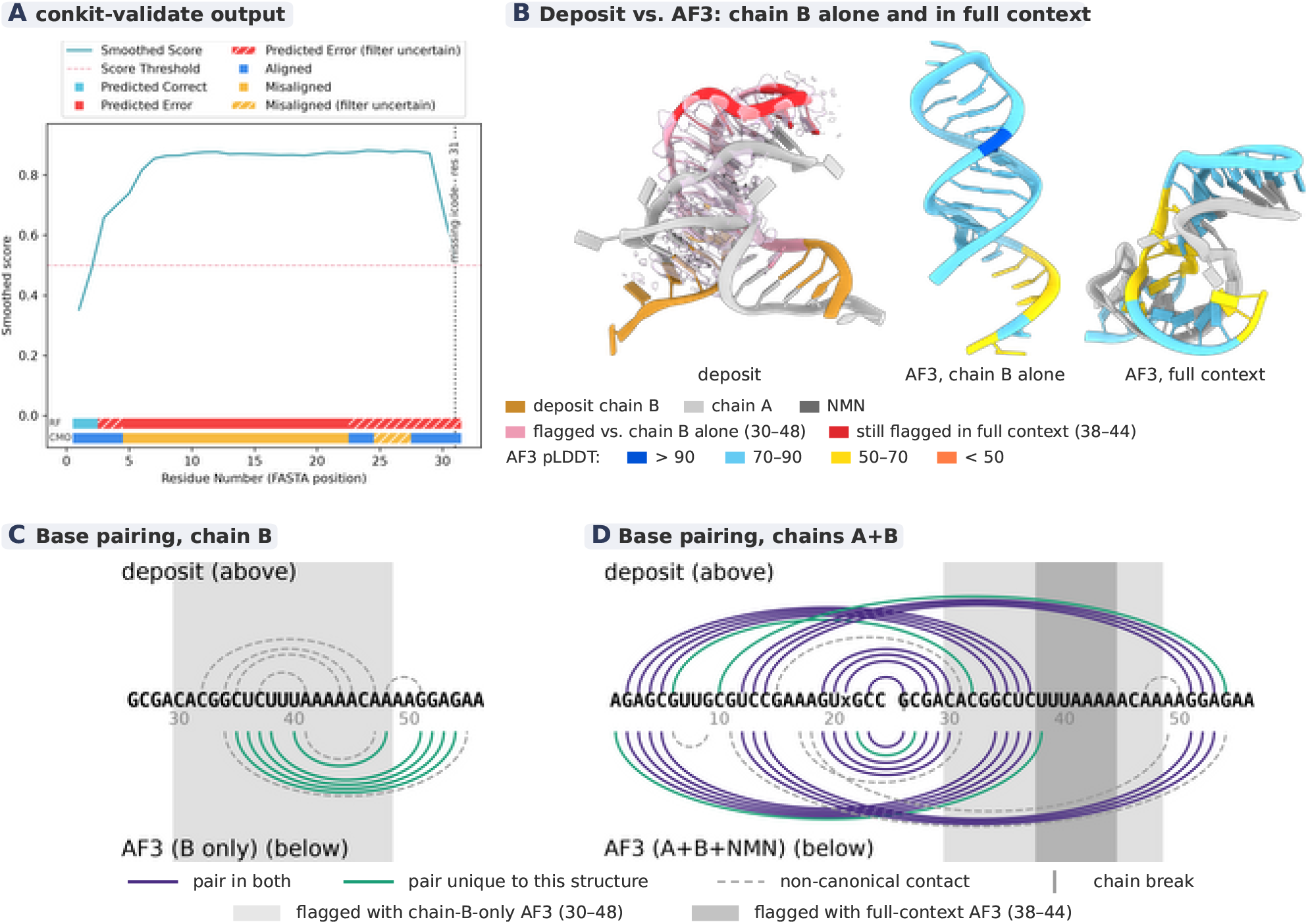
Case study figure for 8I3Z chain B. A: *conkit-validate*’s output plot (smoothed score, RF and CMO tracks) for chain B. B: the deposit and the two AF3 predictions side by side. On B’s left: the deposit, with chain B in ochre, chain A in light grey and the bound NMN in dark grey, in the density map (pale lilac surface, shown within 2.6 Å of chain B). In chain B, the window flagged against the chain-B-only prediction (30-48) is rose, and the part still flagged against the full-context prediction (38-44) red. In B’s middle: the AF3 prediction of chain B alone coloured by per-residue pLDDT. On B’s Right: the full-context AF3 prediction (chains A and B co-folded with NMN), chain A in light grey, predicted chain B is coloured by per-residue pLDDT. C and D: arc diagrams of base pairing, drawn as in Fig. 10 (one key for both, below). C: chain B alone in the deposit (above) and in the AF3 prediction of chain B on its own (below). D: both strands on one axis (chain A 1-24, chain B 26-56, chain break marked at 25) in the deposit (above) and in the full-context AF3 prediction (below). The window flagged against the chain-B-only prediction (30-48) is shown light grey; The window still flagged against the full-context prediction (38-44) is coloured dark grey. Base triples are only partly captured, since each base is assigned at most one partner.

Notably, the map-based CMS method did not flag this high-resolution structure as containing a putative error.

Again, an alternative hypothesis was constructed using *Coot*’s `mutate_base` to match *conkit*’s exact register call. Table 3 shows the resulting ΔRSCC.

**Table 3.** Change in real-space correlation coefficient over the flagged window of 8I3Z chain B (residues 30–48, computed as one region), relative to the raw, unrefined deposited structure (RSCC 0.498), for the deposited baseline and *conkit* ‘s suggested correction. As in Table 1, the deposited baseline’s own change (row 1) is the effect of the refinement pipeline alone. Hypothesis 2’s drop is more than ten times the baseline’s own, the clearest rejection of any case tested.

| hypothesis | ΔRSCC |
|---|---|
| 1: deposited baseline | −0.002 |
| 2: *conkit*’s CMO call (mutate_base) | −0.025 |

Much more clearly than with any case before, the deposited register won out over the one *conkit* suggests, indicating this call is likely a false positive. As interactions between chain A and B are important structural factors in this riboswitch, a second predicted structure was obtained. This time AF3 was provided with all sequence information present in the deposit, making the prediction one of chains A and B co-folded in the presence of the ligand. Rerunning *conkit-validate* using the new, full-context prediction, the size of the flagged error reduced, with only residues 38 to 44 remaining flagged. A structural alignment of the in-context prediction and the deposit shows that they do not share the same global conformation (Fig. 13B).

This case highlights the fact that *conkit* can indeed produce false positives, which are clearly identified based on the map. Important in this regard is the information provided to *conkit*. Here, the validation method failed originally because the predicted structure provided did not match the deposited one. The first prediction lacked the context needed to produce the deposited structure, with chain B folding into a hairpin, while in the deposit it pairs to chain A (Fig. 13B–D). Subsequently, upon being given the necessary context, the size of the flagged error reduced, with the remaining flagged region showing only one pair in the prediction and none in the deposit (Fig. 13D). This remaining flagged region could be attributed either to the difficulty aligning the sequence (UU-UAAAA) it contains, or the globally different conformation (*Gesamt* finds no reasonable alignment between chain B in the deposit and full-context prediction). Additionally chain B has a low number of contacts, overall only 0.26 internal contacts per residue are made, below the 1.45 required by the applicability rule used in the PDB scan, and its *Gesamt* Q-score against the chain-B-only prediction (0.13) falls below the rule’s 0.30 as well.

### Case study from the PDB 9BUQ chain S7: a register error confirmed on independent structural/functional grounds where density comparison could not decide the case

The fifth selected case was that of chain S7 of PDB entry 9BUQ(44), an E-site Met-tRNA in a ribosome of *P. falciparum*. The structure was determined by cryo-EM at a resolution of 2.3 Å. *conkit-validate*’s CMO method indicated a potential register shift error running from residue 17 to residue 70. Differently from before, the predicted size of the register shift wasn’t uniform across this stretch. Rather, it starts at +1 from base 17, then grows to +2 at base 20, and finally grows to +3 from base 32 in a staircase pattern. Such a pattern would be consistent with three deletions, one just before 17, 20 and 32 each. The final +3 shift running through the 3^*′*^ end indicates that the model makes up for these three missing nucleotides by only modelling 74 nucleotides of what is actually a 77-nucleotide molecule. Scores from the RF classifier average 0.846 along this stretch, well above the 0.5 threshold, also indicating it as a putative error, as shown in Fig. 14.

**Fig. 14.**
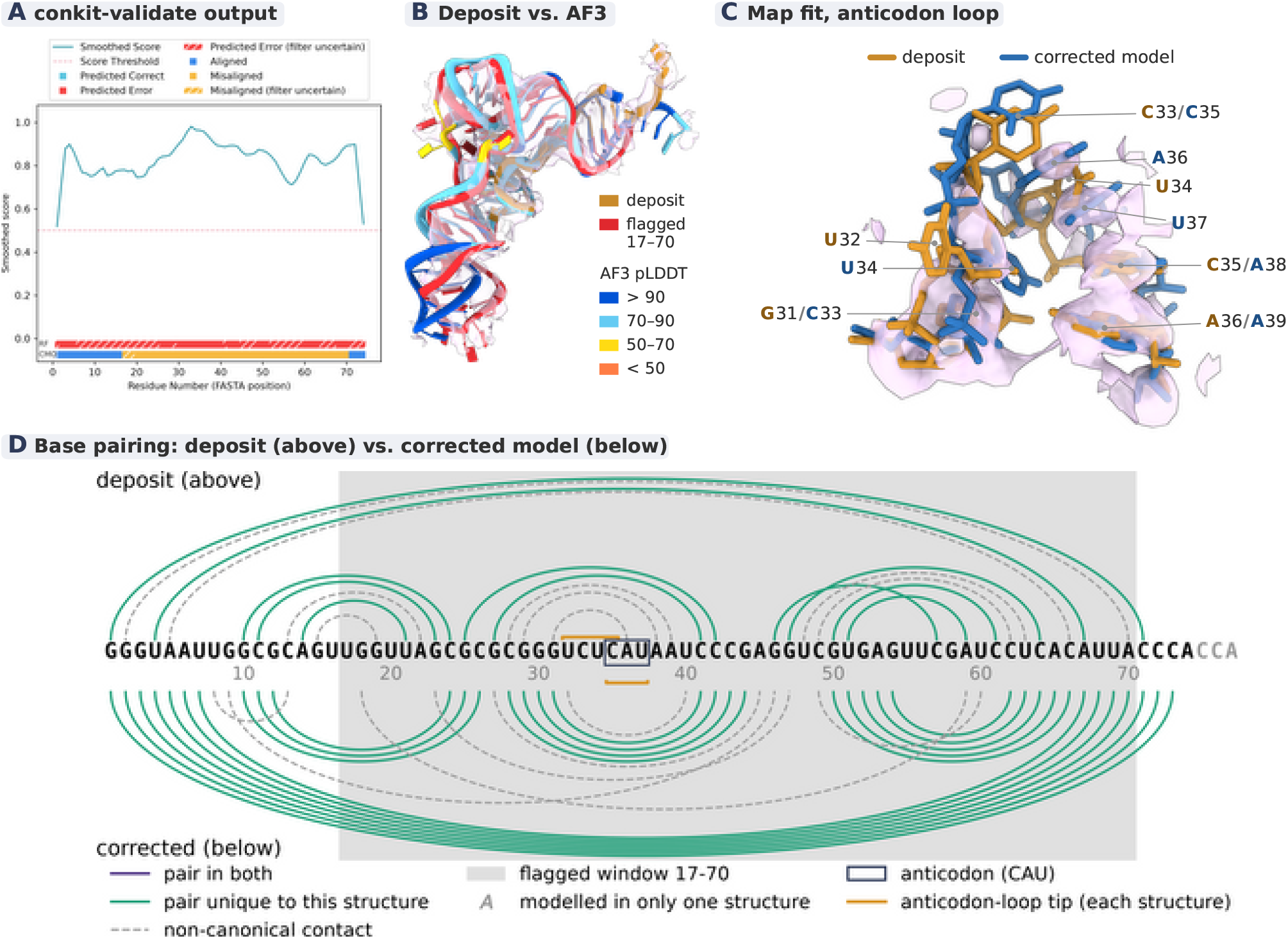
Case study figure for 9BUQ chain S7. A: *conkit-validate*’s output plot (smoothed score, RF and CMO tracks) for chain S7. B: the deposit (ochre, with the putative error in red) superposed with the CCA-appended AF3 prediction (77 nt, coloured by per-residue pLDDT), in the density map (pale lilac surface, shown within 2.6 Å of either model). C: close-up of the anticodon loop of the deposit (ochre, residues 31-36) and the corrected model after refinement (blue, residues 33-39) in the same map. D: arc diagram of the base pairing in the deposit (above the sequence) and the corrected model (below), drawn as in Fig. 10. The appended CCA (grey letters) is not modelled in the deposit; the grey band marks the flagged window (17-70). a navy box marks the Met anticodon (CAU, 35-37), and the amber brackets each structure’s anticodon-loop tip, determined from its base pairing.

Map-based validation by CMS flagged this chain not as having a potential register shift but as potentially mistraced.

Like before, `mutate_base` was used in an attempt to fix the register. The resulting structure was then refined into the map, which resulted in a ΔRSCC of −0.064, making the naive relabel comparable to the raw deposit subjected to the same refinement (yielding ΔRSCC −0.046). Pairing information, however, favoured the AF3 predicted structure, with 19 canonical pairs formed in the corrected model after refinement, compared to 11 in the deposit (see Fig. 14D). Visual inspection of the predicted structure revealed it placed the final three bases (CCA) not as the universal tRNA overhang, even though the CCA overhang was clearly present in the deposited map. Rather, AF3 incorporated C72 and C73 into the acceptor stem (paired to G2 and G1), and placed A74 at the location of the discriminator. A sequence search in GtRNAdb(45) revealed that the sequence in the deposit corresponds exactly to the genomic sequence. However, in many organisms, including *P. falciparum*, the universal CCA overhang on tRNAs is added enzymatically after transcription(46). Such CCA-adding enzymes have been shown to still work on tRNAs with a genomic sequence ending in CCA when the final two cytosines formed part of the acceptor stem and the adenosine took the place of the discriminator(46). The addition of this terminal CCA allows for the good pairing of the AF3 prediction, the presence of the biologically essential and well-resolved CCA overhang, and adherence to *conkit*’s called register shift.

Because of this, a new AF3 prediction was made for the structure of the deposited sequence with the additional, terminal CCA appended, resulting in a structure of 77 rather than 74 nucleotides. Fig. 14B shows the raw deposit and the new, CCA-appended prediction superposed. Unlike in the previous cases, the correction could not be made by relabelling the deposited bases, as it adds three nucleotides; instead, the corrected model was built from the CCA-appended prediction. As the predicted structure had slight conformational differences to the deposit, the new structure was first pulled into the same conformation by matching bases based on *conkit*’s call, so that refinement started from the deposit’s fit to the map rather than from the predicted conformation.Then the restraints pulling our alternative hypothesis into place were relaxed, and the map weight was gradually ramped up during *ISOLDE*’s molecular dynamics flexible fitting (MDFF)(40), the anticodon loop of the result is shown in the map in Fig. 14C. This finally resulted in an RSCC of 0.701, improving over the untouched structure as deposited (0.638) by a ΔRSCC of 0.062. Running the exact deposited structure through the final stages of the same pipeline however produced an RSCC of 0.707 (ΔRSCC of 0.069). The effect of MDFF refinement was thus much greater on the result than that of changing the underlying structure, with the map being unable to differentiate between the 77- and 74-nucleotide models.

Closer study of the deposited structure, however, revealed that at the anticodon-loop tip it exposed the bases UCUC (residues 32-35) rather than the correct Met anticodon CAU, with the cytosine and adenosine of the real anticodon turned into the stem instead of flipped out (Fig. 14C, D). Both AF3 predictions placed the anticodon correctly, folding it into the loop tip as expected of a functional tRNA, and the corrected model keeps it there after refinement (Fig. 14C, D). As the identity of these bases is essential for the tRNA’s function, the register shift error flagged by *conkit* is likely real.

While *conkit-validate* correctly flagged the anticodon region (residues 35, 36 and 37) as having a shifted register, this argument on functional grounds points to the shift being +2 rather than +3 in this region. This is consistent with the third deletion not occurring at position 32 but just after the anticodon (near 38). The benchmark (Fig. 6) already showed that such imprecisions can occur in *conkit*’s suggested fix.

In this case, even with a high experimental resolution (a headline resolution of 2.3 Å and 2.61 Å locally for the chain), no alternative register assignment produced a better model map-fit than the deposit. Nonetheless, *conkit*’s error call could be confirmed. Deeper research into the potential origins of this error revealed two earlier structures of the same molecule, both carrying a tRNA chain with identical sequence to the deposited chain S7 studied here: PDB entry 3J7A(47) (3.2 Å, deposited 2014) and, at lower resolution, PDB entry 3JBP(48) (6.7 Å, deposited 2015). It is likely the register shift error originated there, with those depositors missing the enzymatically appended CCA and thus stretching the genomic CCA end into the overhang. A brief sequence search for PDB entries sharing the same tRNA molecule found the same underlying failure recurring. Different PDB entries associated with the same publication (9BUS, 9BUP and 8TPU(44)), 6OKK, a 2019 entry from the group behind 3J7A(47), and later entries by different groups (9J9H and 9J9I(49), and 9YXB(50)), are all flagged by *conkit* as containing a register error running up to +3, although the specific register reassignments differ. Crucially, no structure has the final CCA appended, while multiple cases fail to expose the correct CAU anticodon.

## Conclusions

The work above shows that, following the modifications made, the contact-prediction-based validation of *conkit-validate* can be extended to RNA structures. In particular, the CMO method works as an effective error-detection technique, as demonstrated on the benchmarking set of artificial data. Notably, however, the success rate of the suggested register reassignment on this data set was worse than in the protein case. This could be either due to the artificial nature of the errors, or because the contact pattern of RNA is primarily based on pairing, providing less redundancy than is present in contact maps from proteins. Parallel to the CMO method, a random forest classifier was trained with the aim of detecting sequence-assignment-related errors that involve more complex reassignments than *map_align* can produce. This classifier has an F1-score of 0.85 (on AF3 structures) slightly below its protein equivalent (F1 0.87(9)), which could be an effect of the artificial nature of its training set. While the RF classifier thus scores better on the artificial set than the CMO method (0.78 on AF3 structures), its calls hold up less well in the PDB scan after filtering for applicability based on structural alignment and number of contacts.

Exactly how well *conkit-validate* works depends on the quality of the provided predicted structure. General prediction models like AF3 work decently, but specialised RNA structure prediction methods like *RNAPro* outperform them. This is encouraging: as RNA structure prediction methods continue to develop, validation by the presented method will become more reliable. Additionally, as seen in the case study concerning 8I3Z, the validation method can be sensitive to large conformational differences. Such cases can be recognised before a call is trusted by checking the global structural alignment (e.g. by *Gesamt* Q-score) and the number of contacts per nucleotide. In the PDB scan above, doing so removed more than half of the putative errors identified by CMO. The development of structural ensemble prediction tools(51), which are being extended to RNA(52), might aid in this.

In light of the continuing development of co-folding methods, which predict the structures of bio-macromolecular complexes (protein, RNA, DNA, …), the expansion of *conkitvalidate* to RNA opens the door to further extending it to multichain validation. This would enable *conkit-validate* to leverage the coevolutionary information present at the interfaces in multichain RNA assemblies, but also protein-RNA and even protein-protein complexes, increasing the applicability to regions with a low number of intra-chain contacts but many inter-chain contacts. Such co-folding information could also aid with issues arising from conformational differences. For example, predicting riboswitches correctly in the holo or apo states could reduce false positives originating in mismatches between the predicted conformation and that observed in experiment. Likewise, general improvement in the quality of RNA structure prediction methods might eliminate systematic mispredictions such as that of the 5S rRNA stem-loop, and thus systematic false positives.

Altogether, this work enables the validation of RNA structures based on coevolutionary information by leveraging structure prediction methods. Such a mapless validation approach is expected to be notably useful in cases of lowresolution density.Because its detection rate does not depend on the local resolution, it remains applicable where map-based methods are not. However, it also proved useful in high-resolution cases where a biologically verifiable error was detected but model-map fit could not differentiate between the error and a proposed correction. As the quality of the error detection is tied to the prediction provided, *conkitvalidate*’s accuracy is expected to improve naturally as prediction methods continue to be developed.

## Data availability

*Conkit-validate* is available as part of the *conkit* Python package which can be found on GitHub at https://github.com/rigdenlab/conkit. As of version 0.14.0 the package is compatible with RNA, that version also contains the trained classifiers presented. *Conkit* is additionally distributed as part of the CCP4 software suite(1), and *conkit-validate* has been integrated into the web-interface*Moorhen* https://moorhen.org/.

## Acknowledgements

AV would like to acknowledge Luc Elliott for advice on automated solving of cryo-EM maps, Eugene Krissinel and Ronan Keegan for help with and access to the *Gesamt* source code, Grzegorz Chojnowski for advice on which structures in the PDB to scan, Emily Johnson and Anthony Evans for advice on machine learning, and CCP4 for funding through a BBSRC grant.

