## supplemental text tables and figures for "A validation approach for RNA structures based on deep-learning 3D-structure predictions"

### Supplementary information

**A. Contact definition and CMO performance.** This section gives the detailed scores behind the choice of contact cut-off (Fig. S1), the confusion matrices and scores of CMO for each prediction input (Fig. S2, table S1), and the share of correctly identified register errors for which CMO also suggests the correct register (table S2).

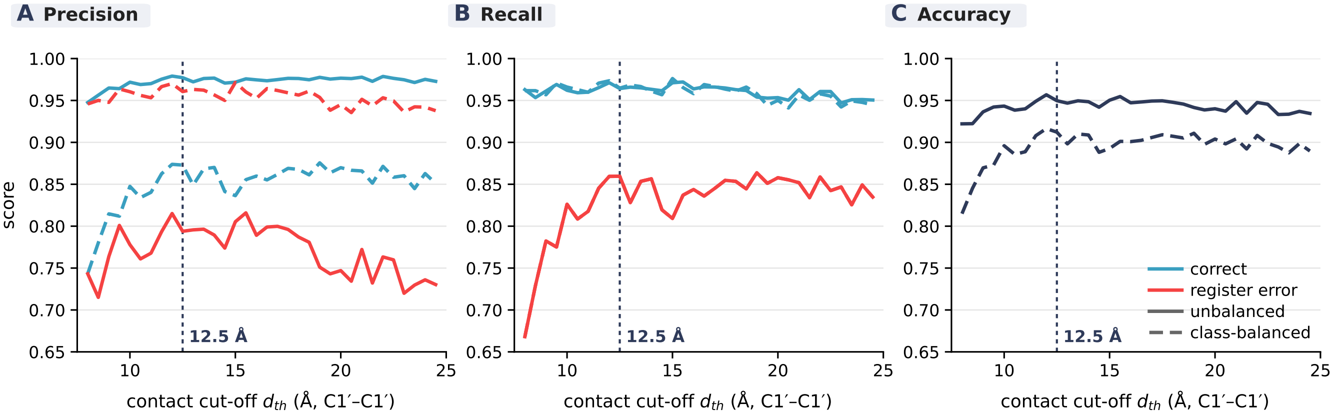


**Fig. S1.** Precision (A), recall (B) and accuracy (C) of CMO as a classifier of register errors, as a function of the contact cut-off $d_{th}$ (C1′-C1′ distance), complementing the F1 scores of Fig. 3. Colours give the class (register errors red, correctly modelled bases blue; accuracy, over both classes, navy), line style the data set: unbalanced (solid) or class-balanced (dashed). The dotted line marks the cut-off used, $12.5$ Å.

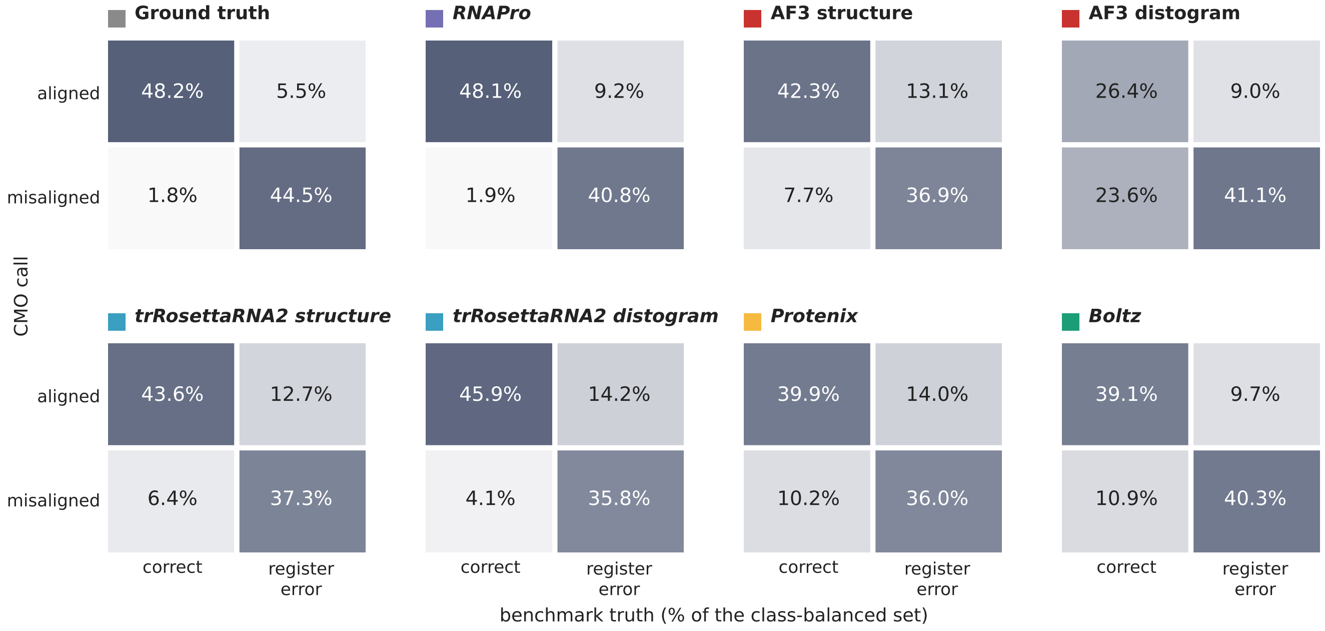


**Fig. S2.** Confusion matrices showing the performance of the CMO approach on RNA using different prediction methods across a version of the benchmark data set balanced to have equal numbers of residues which are part of register errors and correctly modelled. All values are given in percentages of the balanced set, rows give CMO's call for each base (aligned or misaligned), columns the benchmark truth (correctly modelled or part of a register error). Swatches give each input's colour in Fig. 5. No filters were applied to none of the results to allow a fair comparison of input methods.

| input | F1 | accuracy | precision | recall |
|---|---|---|---|---|
| Ground truth | 0.92 | 0.93 | 0.96 | 0.89 |
| *RNAPro* | 0.88 | 0.89 | 0.95 | 0.82 |
| AF3 structure | 0.78 | 0.79 | 0.83 | 0.74 |
| AF3 distogram | 0.72 | 0.67 | 0.64 | 0.82 |
| *trRosettaRNA2* structure | 0.80 | 0.81 | 0.85 | 0.75 |
| *trRosettaRNA2* distogram | 0.80 | 0.82 | 0.90 | 0.72 |
| *Protenix* | 0.75 | 0.76 | 0.78 | 0.72 |
| *Boltz* | 0.80 | 0.79 | 0.79 | 0.81 |

**Table S1.** Performance of CMO as a classifier of register errors using different prediction methods as the input. All calculated for register errors being the positive class on a class balanced version of the benchmark data set.

| input | register correct (%) |
|---|---|
| Ground truth | 76 |
| *RNAPro* | 63 |
| AF3 structure | 46 |
| AF3 distogram | 21 |
| *trRosettaRNA2* structure | 60 |
| *trRosettaRNA2* distogram | 57 |
| *Protenix* | 39 |
| *Boltz* | 38 |

**Table S2.** Percentage of correctly identified register errors where the register suggested by CMO also matched the ground truth register. Results are given per prediction method.

**B. Machine learning classifiers.** This section gives the feature correlations used for the feature reduction (Fig. S3) and the features selected in this way (table S3), followed by the performance of all 24 classifiers (table S4), their ROC curves (Fig. S4) and the feature importances from the ablation study (Fig. S5).

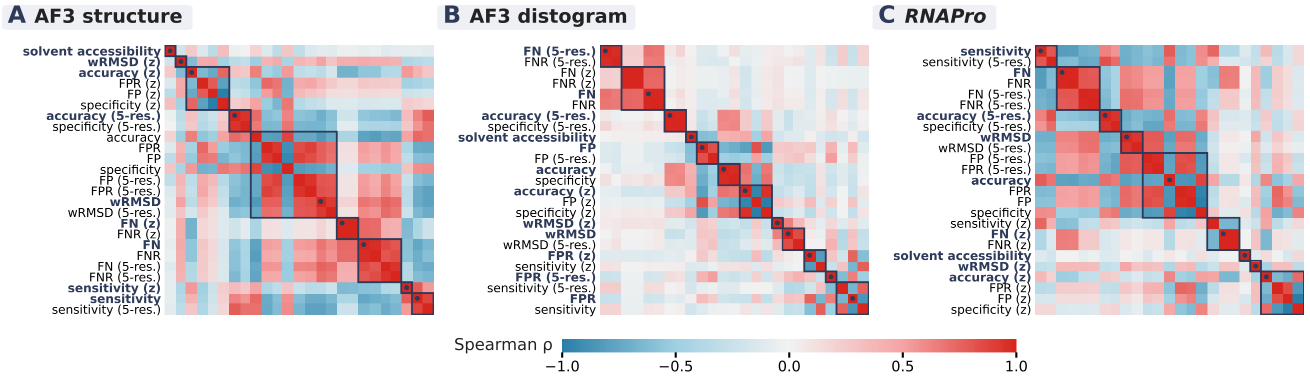


**Fig. S3.** Spearman correlation between the 25 *conkit* features over the training set, for the feature-reduction clustering per prediction method: (A) AF3 structures, (B) AF3 distograms and (C) *RNAPro* structures, without CANA codes. Features are ordered by the clustering; boxes on the diagonal mark the clusters (Ward linkage on $1-|\rho|$, cut at $0.4$), and the feature kept from each cluster is shown in bold with a dot. (z): z-score relative to the bases within $20\,\text{Å}$; (5-res.): averaged over five consecutive residues.

| **Prediction method** | **including CANA** | **without CANA** |
|---|---|---|
| AF3 structures | ACC, AAA, AAw, AAu, A-B, B-A, BBB, BBw, B12, BB2, miB, ICL, OPN, SYN, ZZZ, WRMSD, FN, SENSITIVITY, ACCURACY SMOOTH, ZSCORE WRMSD, ZSCORE ACCURACY, ZSCORE FN, ZSCORE SENSITIVITY | ACC, WRMSD, FN, SENSITIVITY, ACCURACY SMOOTH, ZSCORE WRMSD, ZSCORE ACCURACY, ZSCORE FN, ZSCORE SENSITIVITY |
| AF3 distograms | ACC, AAA, AAw, AAu, A-B, B-A, BBB, BBw, B12, BB2, miB, ICL, OPN, SYN, ZZZ, WRMSD, ACCURACY, FN, FP, FPR, ACCURACY SMOOTH, FN SMOOTH, FPR SMOOTH, ZSCORE WRMSD, ZSCORE ACCURACY, ZSCORE FPR | ACC, WRMSD, ACCURACY, FN, FP, FPR, ACCURACY SMOOTH, FN SMOOTH, FPR SMOOTH, ZSCORE WRMSD, ZSCORE ACCURACY, ZSCORE FPR |
| *RNAPro* | ACC, AAA, AAw, AAu, A-B, B-A, BBB, BBw, B12, BB2, miB, ICL, OPN, SYN, ZZZ, WRMSD, ACCURACY, FN, SENSITIVITY, ACCURACY SMOOTH, ZSCORE WRMSD, ZSCORE ACCURACY, ZSCORE FN | ACC, WRMSD, ACCURACY, FN, SENSITIVITY, ACCURACY SMOOTH, ZSCORE WRMSD, ZSCORE ACCURACY, ZSCORE FN |

**Table S3.** Features selected by correlation-clustering reduction, organised by input prediction method (*RNAPro* structures, AF3 structures and AF3 distograms), for both the case where CANA codes were available as features and where they were left out.

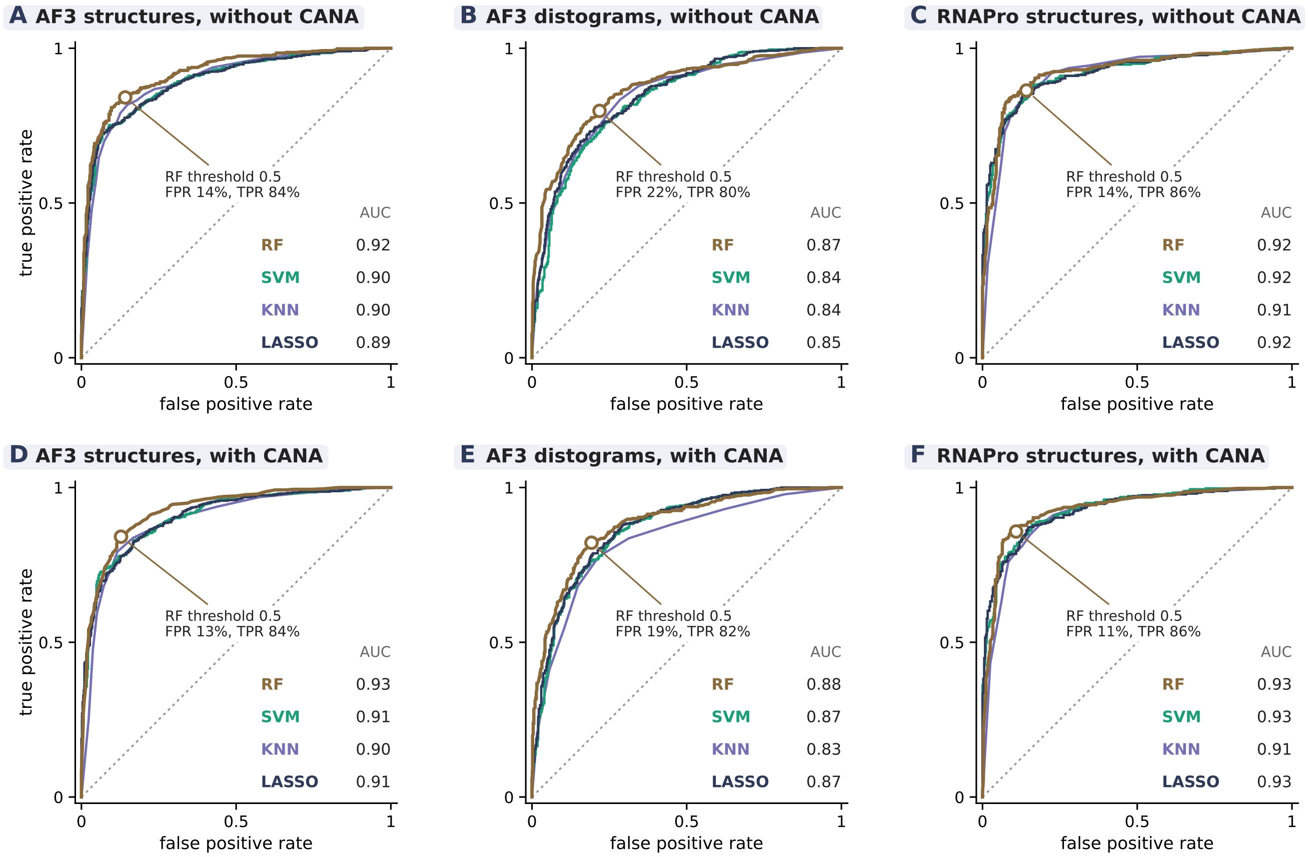


**Fig. S4.** ROC curves of the four ML classifiers (RF, SVM, KNN, LASSO) on class-balanced validation sets, per prediction input (AF3 structures, AF3 distograms, *RNAPro* structures), (A,B,C) without and (D,E,F) with CANA codes. The open circle marks the RF classifier at its default threshold of $0.5$, as used by *conkit-validate*. Random assignment is shown by the grey dotted line.

| score | F1 | ROC-AUC | accuracy | precision | recall |
|---|---|---|---|---|---|
| Method | AF3 structures with CANA codes | | | | |
| RF | 0.85 | 0.93 | 0.86 | 0.87 | 0.84 |
| SVM | 0.82 | 0.91 | 0.82 | 0.83 | 0.81 |
| KNN | 0.84 | 0.90 | 0.84 | 0.83 | 0.84 |
| LASSO | 0.82 | 0.91 | 0.82 | 0.83 | 0.81 |
| Method | AF3 structures without CANA codes | | | | |
| RF | 0.85 | 0.92 | 0.85 | 0.86 | 0.84 |
| SVM | 0.81 | 0.90 | 0.81 | 0.81 | 0.81 |
| KNN | 0.83 | 0.90 | 0.83 | 0.85 | 0.82 |
| LASSO | 0.81 | 0.89 | 0.81 | 0.82 | 0.81 |
| Method | AF3 distograms with CANA codes | | | | |
| RF | 0.82 | 0.88 | 0.81 | 0.81 | 0.82 |
| SVM | 0.79 | 0.87 | 0.78 | 0.78 | 0.80 |
| KNN | 0.78 | 0.83 | 0.78 | 0.78 | 0.78 |
| LASSO | 0.79 | 0.87 | 0.79 | 0.80 | 0.78 |
| Method | AF3 distograms without CANA codes | | | | |
| RF | 0.79 | 0.87 | 0.79 | 0.78 | 0.80 |
| SVM | 0.76 | 0.84 | 0.76 | 0.75 | 0.78 |
| KNN | 0.77 | 0.84 | 0.77 | 0.76 | 0.78 |
| LASSO | 0.76 | 0.85 | 0.77 | 0.78 | 0.75 |
| Method | *RNAPro* structures with CANA codes | | | | |
| RF | 0.87 | 0.93 | 0.87 | 0.89 | 0.86 |
| SVM | 0.85 | 0.93 | 0.85 | 0.88 | 0.81 |
| KNN | 0.85 | 0.91 | 0.85 | 0.85 | 0.85 |
| LASSO | 0.84 | 0.93 | 0.85 | 0.89 | 0.78 |
| Method | *RNAPro* structures without CANA codes | | | | |
| RF | 0.86 | 0.92 | 0.86 | 0.86 | 0.86 |
| SVM | 0.84 | 0.92 | 0.85 | 0.88 | 0.81 |
| KNN | 0.85 | 0.91 | 0.85 | 0.87 | 0.83 |
| LASSO | 0.84 | 0.92 | 0.84 | 0.88 | 0.79 |

**Table S4.** Performance (F1, ROC-AUC, accuracy, precision, recall) of the four ML classifiers (RF, SVM, KNN, LASSO) across prediction methods and with/without CANA codes on class-balanced validation sets. All metrics except the ROC-AUC use the default classifier score threshold of $0.5$.

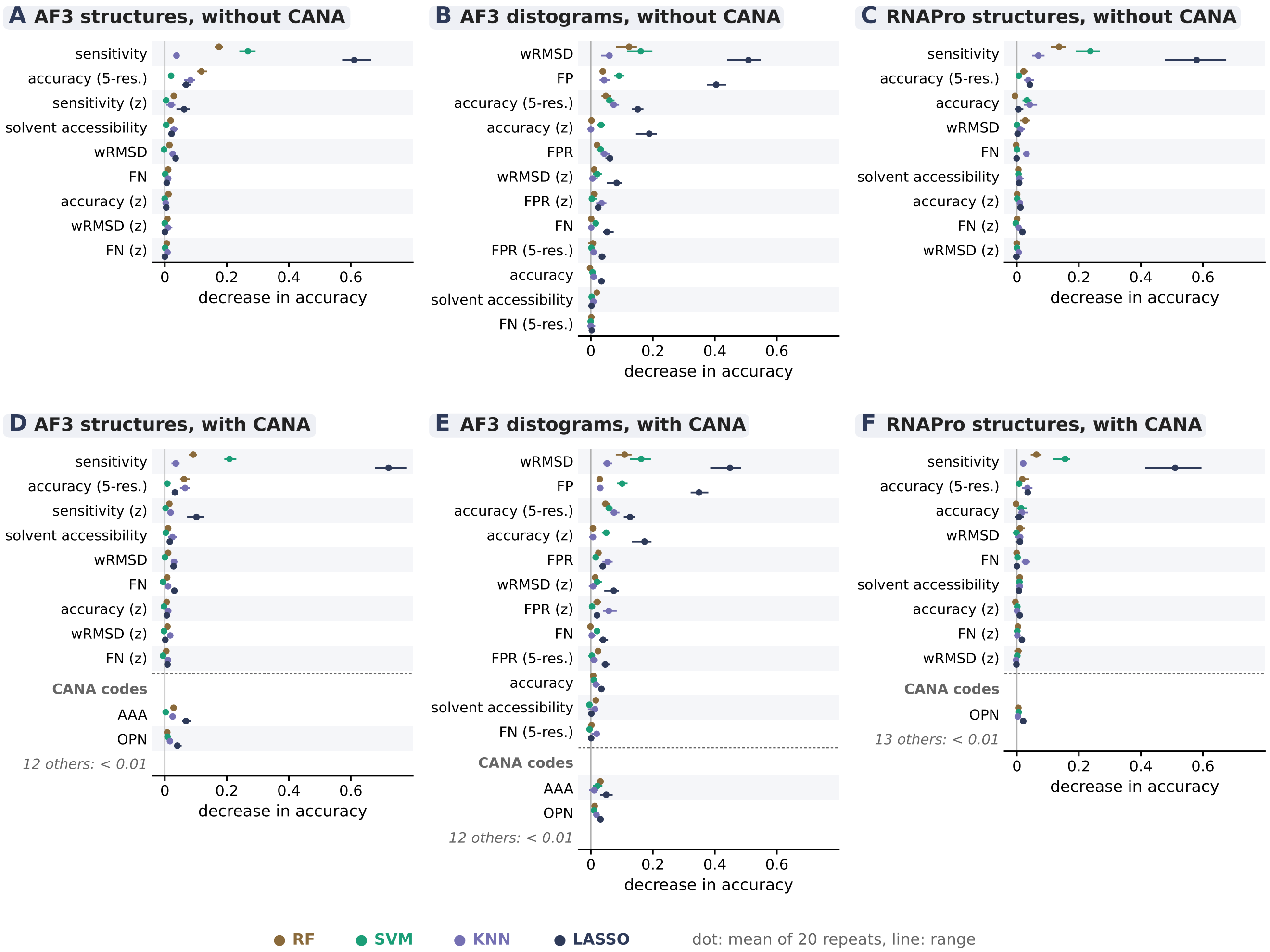


**Fig. S5.** Feature importance for the four ML classifiers per prediction input, (A,B,C) without and (D,E,F) with CANA codes: the decrease in validation accuracy when the values of one feature are randomly permuted, repeated 20 times (dot: mean, line: range). Features are ordered by their mean importance over the four algorithms without CANA codes; names as in Fig. S3. CANA-code features with an importance below $0.01$ for every algorithm are not shown individually.

**C. Additional filtering of the PDB-wide scan.** The per-residue false-positive filter (Methods, Filtering) was trained on artificial errors, which contain no unfolded deposits and few global conformational differences between prediction and model. Therefore chains where *conkit-validate* struggles to apply as a whole, are not filtered out. To address this a chain-level applicability rule was created based on two properties: the global *Gesamt* Q-score between the deposited chain and the prediction, and the number of contacts per residue in the deposited chain. For *Conkit-validate* (CMO and RF) to be considered applicable to a chain both need to be above a threshold, Q $\geq 0.30$ and at least $1.45$ contacts per residue (Fig. S6).

The two thresholds were determined using an active learning approach on the set of 695 chains flagged by CMO or RF. Through visual inspection, chains were labelled "not applicable" when they were (partially)unfolded, or when a globally different conformation was predicted (as for 8I3Z, Fig. 13). Notably the accuracy of calls was not taken into account, that means calls where *conkit-validate* is applicable but wrong are in the "applicable" class, making the created rule purely applicability directed, rather than dealing with false-positives as a whole. Labelling happened blind to the chain's Q-score and contacts.

The active learning protocol started from a Bayesian posterior probability distribution with a learnt label-noise rate, over a grid of threshold pairs, seeded using five chains of known status. The posterior was updated after each label, and the next chain to label was the one whose label was most uncertain under the current posterior. At most one chain per PDB entry was queried, and near-identical (Q, contacts) pairs were down-weighted. The 185 flagged 5S rRNA chains, with nearly identical Q-scores, were counted as a single data point: "applicable" when they differed from the prediction only in the one stem-loop that AF3 systematically predicts differently. Training stopped after 59 labels, when the most uncertain remaining chain had an uncertainty below $0.1$ (posterior in Fig. S6B).

The created applicability rule with set thresholds was then tested on a random sample of 36 chains (ensuring no overlap with the training data), stratified over the four quadrants created by the thresholds (Fig. S6A, filled markers). In this set 20 chains passed the learnt applicability rule and 2 of those were visually judged not applicable, and none of the 16 chains it excludes was judged applicable. Across all chains in the PDB-scan, 4020 of the 7556 (with both values computed) are applicable by this rule. Of the flagged chains CMO keeps 255 of 565 and RF 31 of 299.

During active learning it was noted many 5S rRNA had visually good alignments but had a common false positive in a smaller mispredicted stem-loop. For training the rule these were kept as "applicable", as any error flagged in most of the chain would not be attributed to low contacts or poor conformational agreement. However, almost all 5S rRNA chains with Q $< 0.45$ (having the common misprediction) (537 chains, 233 of them applicable) are eukaryotic, about $85\%$ mammalian among those flagged by CMO. All six of the flagged chains that were inspected were false positives in that stem-loop, and CMS flags only 1 of the 124. As they pass the applicability rule, they are not removed, but shown separately (dotted line in Fig. 9).

For CMS, the resolution range was re-judged per chain for EM structures. One local resolution was determined per chain: the mask-corrected masked half-map FSC $= 0.143$ resolution, computed from the deposited half maps cropped around the chain, with a soft mask of 3 Å around the chain's atoms and phase-randomisation correction as in *RELION* post-processing. It could be measured for 3059 of the 4713 EM chains, with the others missing deposited half maps, lying outside the box, or computation of the resolution failing due to a variety of issues (half maps not being independent, a value appearing at the box limit or a local resolution $> 8$ Å worse than the headline resolution). The results of CMS are only counted for chains in the stated 5 Å range of applicability, judged on the per chain local resolution where available. For chains without a measurable local resolution the headline resolution was used, or they were left out in figures S7 and S8.

Many CMS register-shift calls not corroborated by *conkit* are tRNAs in large ribosome structures. They tend to lie in significantly worse local density than unflagged tRNAs in the same maps (paired comparison over 50 of 75 entries, Wilcoxon $p = 10^{-4}$).

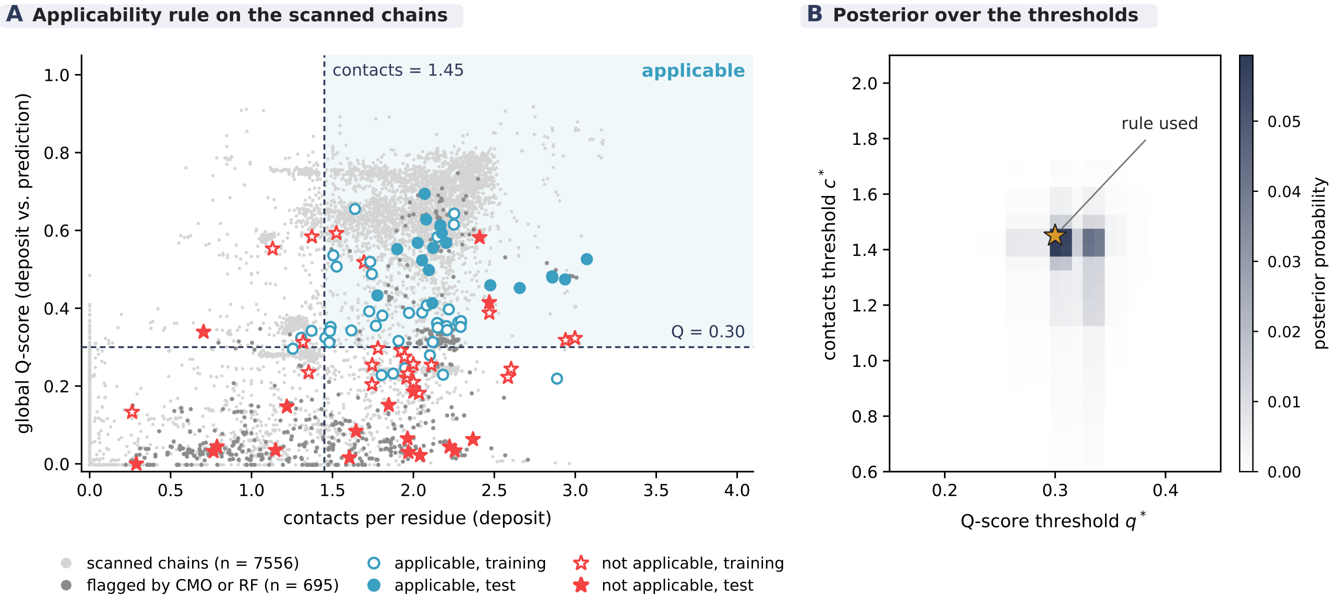


**Fig. S6.** Applicability rule for *conkit-validate* applied in the PDB-scan. (A) Global *Gesamt* Q-score between deposit and AF3 prediction against the number of contacts per residue in the deposit, for the 7556 scanned chains with both values (light grey), the 695 chains flagged by CMO or RF (dark grey), and the chains labelled as applicable (blue circles) or not applicable (red stars) during training (open markers, including the five seeds) and testing (filled markers). Dashed lines: the thresholds of the rule. shaded: the region where *conkit-validate* is considered applicable. (B) Posterior probability over the two thresholds after training, with the thresholds used (star).

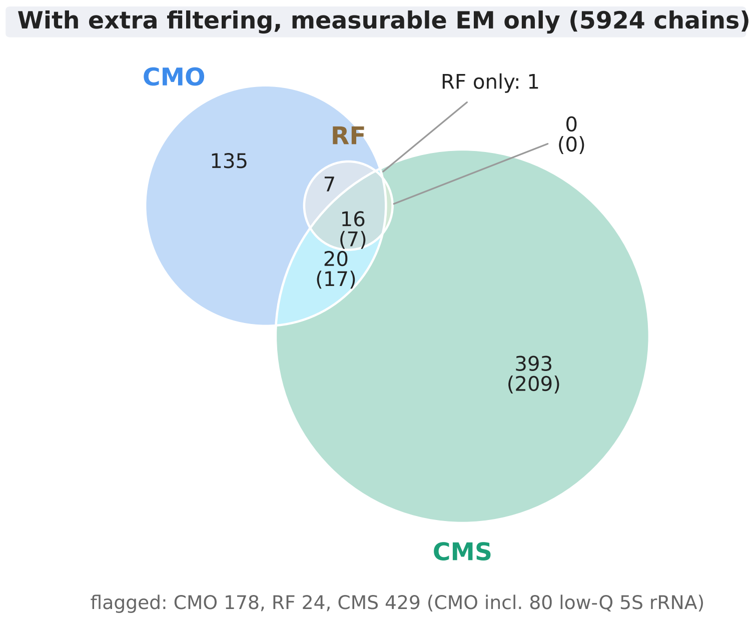


**Fig. S7.** Chains flagged as containing putative errors by *conkit-validate* using the CMO (blue) and the random forest classifier (RF, brown), and by *checkMySequence* (CMS, green, any error type), out of the 5924 chains validated by *conkit-validate* that remain when EM chains without a measurable local resolution are left out, and with the additional filters described in Supplementary Section C: CMO and RF flags are only counted for chains passing the applicability rule (global Q-score $\geq 0.30$ and $\geq 1.45$ contacts per residue), CMS flags only for chains within its $5\,\text{Å}$ working range, judged for EM chains by the local resolution of the chain itself (83 chains removed). Of the 178 filtered CMO chains, 80 are low-Q 5S rRNA chains (Supplementary Section C). The bracketed number in every region CMS appears in gives how many of its chains were flagged specifically for a register shift. The full version, with those EM chains on their headline resolution, is Fig. 8B.

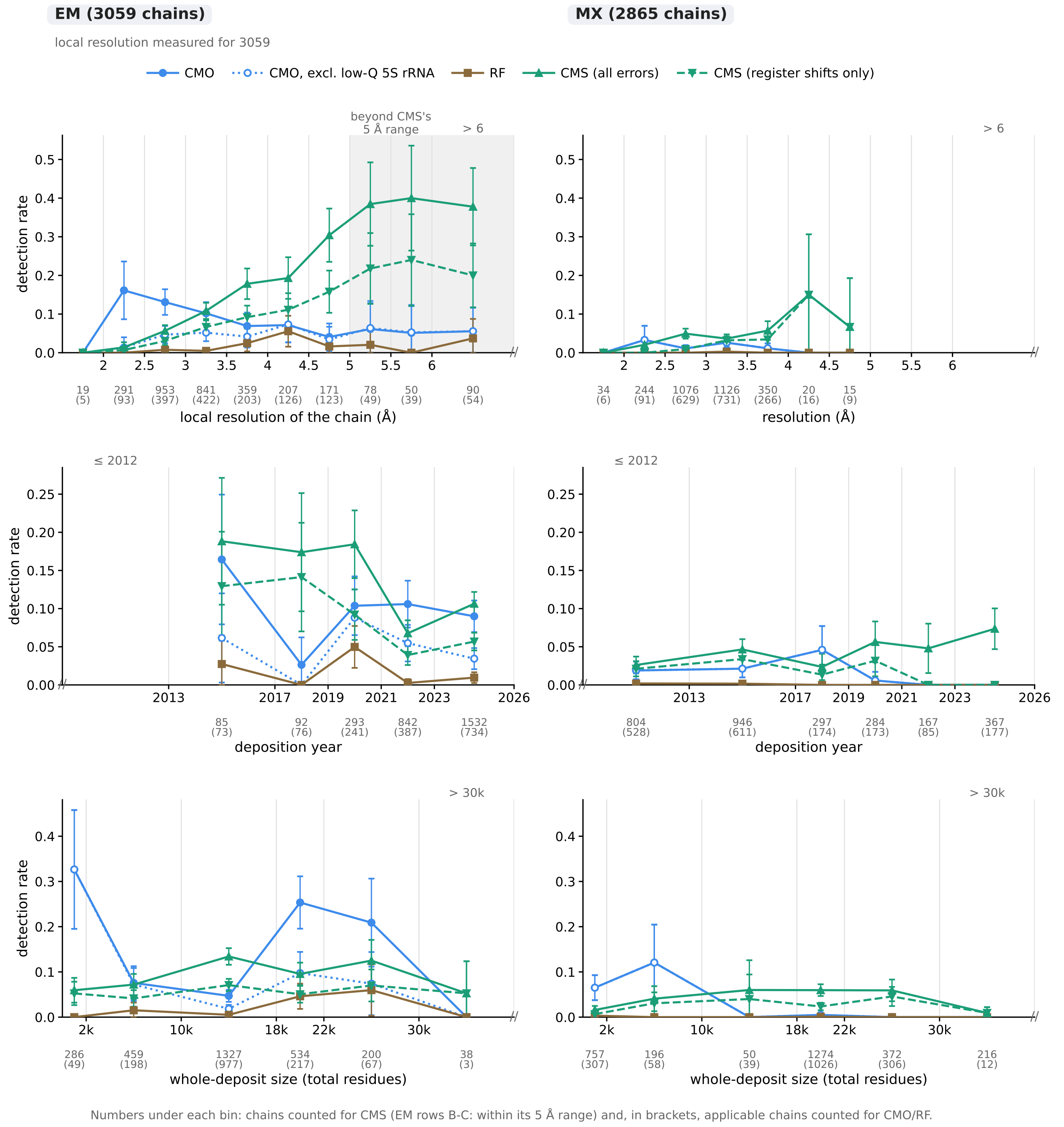


**Fig. S8.** Detection rate of putative errors by CMO (blue, solid; dotted: excluding low-Q 5S rRNA chains), RF (brown), CMS (green) and CMS considering only register-shift errors (green, dashed), for structures determined by electron microscopy (EM, left) and X-ray crystallography (MX, right), against (A) resolution, (B) deposition year and (C) whole-deposit size (total number of modelled residues in the entry). All panels use the additional filtering of Fig. 8: CMO and RF rates are computed over applicable chains only. For EM, resolution is the local resolution of each chain's own density leaving out all other chains (3059 EM chains remain), in (A) bins beyond CMS's $5\,\text{Å}$ working range are shaded, in (B) and (C) CMS only counts chains within that range. Numbers under each bin: chains counted for CMS and, in brackets, applicable chains counted for CMO/RF; error bars: $95\%$ intervals. The full version, with those EM chains on their headline resolution, is Fig. 9.

**D. Q-scores for 7L48 chain E.** EMD-23157 has no deposited half maps, so the per-chain local resolution used for the PDB-wide scan could not be calculated for 7L48. To check the relative quality of the density around the flagged region, instead per-residue Q-scores(42) were calculated for the deposited model in its map. This was done with *ChimeraX*(21) (version 1.11.1) using the `qscore` command (QScore bundle 1.2 from the *ChimeraX* Toolshed) at its default settings.Two runs gave identical values. Residues of chain E were grouped into the two flagged windows (133 to 140 and 182 to 190) and the rest of the chain. The windows (median Q-score 0.34 and 0.33) lie in weaker density than the rest of chain E (median 0.48; two-sided Mann–Whitney U test on per-residue values, $p = 1.5 \times 10^{-4}$) and than the protein chain (median 0.44).
